# Real-time measurements of ATP dynamics via ATeams in *Plasmodium falciparum* reveal drug-class-specific response patterns

**DOI:** 10.1101/2023.12.20.572598

**Authors:** Eric Springer, Kim C. Heimsch, Stefan Rahlfs, Katja Becker, Jude M. Przyborski

## Abstract

*Malaria tropica*, caused by the parasite *Plasmodium falciparum* (*P. falciparum*) remains one of the greatest public health burdens for humankind. Due to its pivotal role in parasite survival, the energy metabolism of *P. falciparum* is an interesting target for drug design. To this end, analysis of the central metabolite ATP is of great interest. So far, only cell disruptive or intensiometric ATP assays have been available in this system, with various drawbacks for mechanistic interpretation, and partly inconsistent results. To address this, we have established fluorescent probes, based on FRET and known as ATeam, for use in blood stage parasites. ATeams are capable of measuring MgATP^2-^ levels in a ratiometric manner, thereby facilitating *in cellulo* measurements of ATP dynamics in real-time using fluorescence microscopy and plate reader detection, and overcoming many of the obstacles of established ATP analysis methods. Additionally, we established a superfolder variant of the ratiometric pH sensor pHluorin (sfpHluorin) in *P. falciparum* to monitor pH homeostasis and control for pH fluctuations, which may affect ATeam measurements. We characterized recombinant ATeam and sfpHluorin protein *in vitro* and stably integrated the sensors into the genome of the *P. falciparum* NF54*attB* cell line. Using these new tools, we found distinct sensor response patterns caused by several different drug classes. Arylamino alcohols increased and redox-cyclers decreased ATP, doxycycline caused first-cycle cytosol alkalization, and 4-aminoquinolines caused aberrant proteolysis. Our results open up a completely new perspective on drugs’ mode of action with possible implications for target identification and drug development.

## Introduction

Malaria is a serious infectious disease and remains one of the greatest public health burdens for humankind. In 2022, an estimated 249 million cases occurred worldwide. Of these, around 608,000 people died, with the majority of victims being children under five. Of *Plasmodium* species, *P. falciparum* has the highest impact on global health, causing most infections as well as the deadliest form of the disease (WHO 2023). With growing drug resistance, basic research for development of new drugs remains critical for global health.

In the asexual blood stages it was shown, using isotopic labelling (MacRae *et al*. 2013) and extracellular flux analysis (Sakata-Kato and Wirth 2016), that the parasites rely mainly on glycolysis for adenosine triphosphate (ATP) production and use oxidative phosphorylation (OXPHOS) to only a minimal extent. Due to this limited metabolic flexibility, glycolytic enzymes (Jortzik and Becker 2011), as well as upstream processes such as glucose transporters (Jiang 2022) were proposed as potential drug targets. These data also further highlight the importance of ATP as the main energy source for these parasites, and stimulate research into ATP metabolism.

There are various approaches for measuring ATP that are extensively reviewed elsewhere (Ley-Ngardigal and Bertolin 2022). In *P. falciparum*, most ATP measurement approaches to date used luciferase-based bioluminescence assays to measure ATP (Kanaani and Ginsburg 1989, 1992; Fry *et al*. 1990; Kirk *et al*. 1996; Saliba and Kirk 1999; Ayi *et al*. 2009; Khan *et al*. 2012; Lelièvre *et al*. 2012; van Schalkwyk *et al*. 2013; Miguel-Blanco *et al*. 2017; Shi *et al*. 2021; Morais *et al*. 2022; Peatey *et al*. 2012) or transgenic luciferase-expressing cell lines as a surrogate for parasite viability (Khan *et al*. 2012; Cevenini *et al*. 2014). However, these approaches have inherent flaws regarding their applicability for mechanistic drug-response interpretation. ATP-response towards drug interventions in luciferase-based bioluminescence ATP assays is usually compared to the output of a control group and based on the parasite count, as the exact parasitic cytosolic volume at a given time is hard to define. While using estimations of average parasite volume to calculate a definite [ATP] can partially circumvent this issue, this approach is even less applicable when comparing [ATP] between a control and a drug intervention group in situations in which growth differences are expected, as many drug treatments are known to interfere with protein biosynthesis (Sheridan *et al*. 2018). If a given intervention causes growth retardation and a reduction in cell volume, the intensity of a luciferase-based bioluminescence assay reaction normalised to a certain cell count can decrease in the intervention group, even when the actual molar [ATP] are identical. Additionally, measurements cannot easily be applied to subcellular compartments and are prone to influence from sample preparation. While the intensiometric approach using luciferase expressing cell lines can be applied to specific subcellular compartments, it also does not necessarily reflect true differences in ATP levels, as the output is responsive to other factors that are expected to change through drug interventions, such as levels of sensor expression or degradation. These difficulties were highlighted by Khan *et al*. (2012), who used both a cell disruptive luciferase-based bioluminescence assay and a transgenic luciferase-expressing cell line to study ATP-drug response in trophozoite stages. This study showed that while mefloquine and artemisinin caused a clear increase in amount of ATP over 10 hours, the luminescence of the transgenic cell line decreased strongly under the same conditions.

To overcome these limitations, we decided to establish genetically encoded ratiometric FRET-based MgATP^2-^ sensors from the ATeam (Adenosine 5ʹ-Triphosphate indicator based on Epsilon subunit for Analytical Measurements) family in the *P. falciparum* NF54*attB* strain. ATeams were developed by Imamura *et al*. (2009) and offer real-time *in cellulo* measurements of ATP dynamics on a single-cell or even compartmental level. ATeams are composed of a circular permutated mVenus yellow fluorescent protein (YFP) and an mseCFP cyan fluorescent protein (CFP) connected with an ATP binding domain, the ε-subunit of a *Bacillus subtilis* F_o_F_1_-ATP synthase. Upon ATP binding, this domain changes its conformation, which brings YFP and CFP into close proximity and increases the FRET interaction. Analysis of the YFP and CFP specific emission ratio after CFP excitation demonstrates a correlative response towards MgATP^2-^. Thus, this approach should reflect MgATP^2-^ levels independently of the sensor expression level. Please note that the physiological relevant form of ATP is MgATP^2-^, however we shall follow convention and simply refer to this as ATP, unless otherwise required for clarity.

Modification of the ATP binding domain led to a series of ATeams with different sensitivities towards ATP (Imamura *et al*. 2009). Furthermore, red-shifted GO-ATeams were developed based on green fluorescent protein (GFP) and orange fluorescent protein (OFP [Nakano *et al*. 2011]). In addition, the dimerization interface of GFP-based proteins was modified to yield sensors with improved dynamic range. Kotera *et al*. (2010) discovered that the L206A mutation yielded the highest dynamic range. We therefore chose the ATeam1.03-nD/nA sensor with an improved dynamic range and sensitivity in the low millimolar range, as well as the ATeam1.03YEMK mutant with a slightly higher sensitivity towards ATP. Despite their advantages over other ATP-analysis methods, GFP-based genetically-encoded sensors also have pitfalls to consider. Most importantly, the fluorescence of GFP-based proteins may exhibit sensitivity to pH, as reviewed elsewhere (Shaner *et al*. 2005). Treatment with antiparasitic compounds might induce pH changes in *P. falciparum* and it has previously been shown that illumination itself can induce a pH drop (Wissing *et al*. 2002). To control for this, we in parallel established a parasite line expressing a genetically encoded pH sensor, to monitor possible pH changes caused by drug interventions and sample preparation. pH indicator systems, such as pHluorin (Kuhn *et al*. 2007) or Sypher (Rahbari *et al*. 2017) have already been established in *P. falciparum*. Nevertheless, we chose to establish the superfolder variant developed by Reifenrath and Boles (2018). Schuh *et al*. (2018) found that the superfolder mutations appear to cause increased fluorescence intensity in the parasites. Establishing of the super folder variant of pHluorin is therefore likely of advantage with respect to increased signal-to-noise ratio.

Another pitfall to consider is the possibility of drug-sensor interactions which might lead to potential experimental artefacts. We therefore recombinantly produced ATeam and sfpHluorin proteins to analyse drug-sensor interactions *in vitro,* and thereby, control for potential confounding effects.

With these sensors, we aimed to establish a simple and robust platform to analyse ATP-drug response levels and make these tools available for the malaria community. Additionally, we analysed the ATeam response towards exposure to a selection of different antiparasitic compound classes. These include the 4-aminoquinolines chloroquine (CQ), amodiaquine (AQ), and pyronaridine (PYRO), the arylamino alcohols quinine (QN), mefloquine (MQ), and lumefantrine (LUM), dihydroartemisinin (DHA), the redox-cycler methylene blue (MB [Schirmer *et al*. 2011]), and plasmodione (PD [Ehrhardt *et al*. 2016]), the apicoplast-targeting doxycycline (DOXY [Dahl *et al*. 2006]), the electron transport chain (ETC)-targeting atovaquone (ATQ [Fry and Pudney 1992]), the protein synthesis inhibitor cycloheximide (CHX [Geary and Jensen 1983]), as well as the PfGluPho-targeting drug candidate SBI 0797750 (SBI [Berneburg *et al*. 2022]). With these, we wanted to provoke possibly distinct sensor responses to gain new insights into their mode of action.

## Results

### Recombinant ATeam proteins show a concentration-dependent increase in emission ratio towards ATP

To measure potential direct drug-sensor interactions and to characterize ATeam1.03-nD/nA and ATeam1.03YEMK sensors side-by-side for the first time, we produced His-tagged ATeam proteins recombinantly expressed in *E. coli* and purified them using affinity chromatography. Titration of the ATeam1.03-nD/nA and ATeam1.03YEMK proteins with ATP caused a dose dependent emission ratio increase, as reported by Kotera *et al*. (2010) and Imamura *et al*. (2009), respectively (Fig. 1). We saw a higher dynamic range for ATeam1.03-nD/nA and a higher sensitivity towards ATP for ATeam1.03YEMK. We also confirmed that neither sensor showed a response to AMP or ADP, as previously shown by De Col *et al*. (2017) and Imamura *et al*. (2009).

**Fig. 1:**
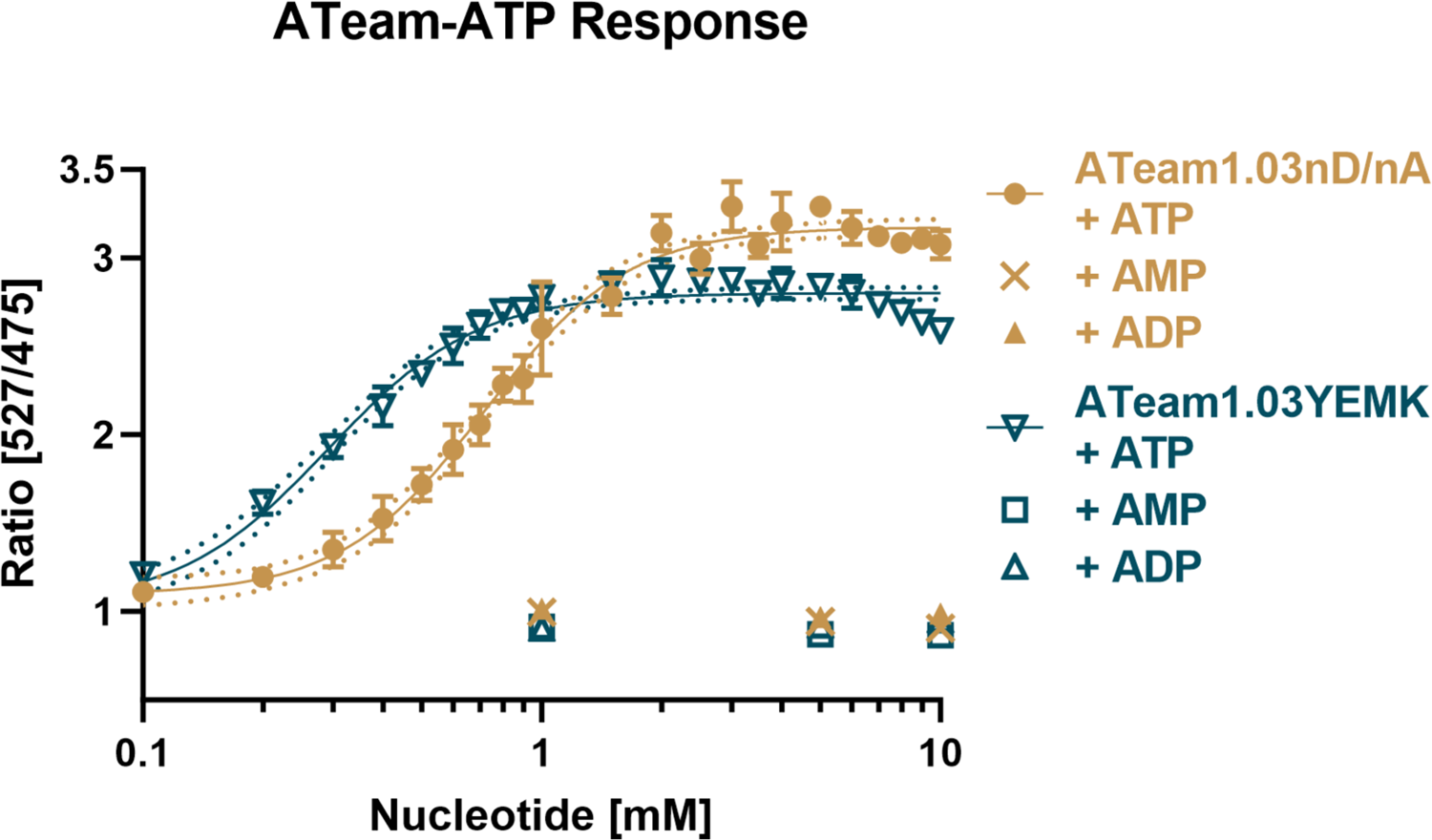
ATeam1.03-nD/nA and ATeam1.03YEMK show a dose-dependent ratio increase in response to ATP. Nucleotides were dissolved in equimolar solution of MgCl_2_. Measurements were conducted in a plate reader via excitation at 435 nm and emission sensing at 527 nm for YFP and 475 nm for CFP. Mean ratio of n = 3 independent experiments is shown. Error bars indicate SD.

We further compared the pH sensitivity, emission spectra and time-responsiveness of both sensors. Despite small discrepancies likely due to different buffer conditions, the ratio response trend is generally in accordance with Imamura *et al*. (2009), Kotera *et al*. (2010), and De Col *et al*. (2017). We saw an apparent increased dynamic range of ATeam1.03-nD/nA towards pH 6 (Fig. 2A). However, the sensor also showed different ratios at different pH even without the presence of ATP, indicating pH sensitivity of the fluorophores. In contrast, the ratio of ATeam1.03YEMK without ATP substrate was almost invariant between pH 6 and 9 (Fig. 2B). ATeam1.03YEMK showed an apparent lower dynamic range from pH 7 to 5. However, for ATeam1.03YEMK, this was accompanied by an increased CFP and a decreased YFP emission (Fig. S1D), corresponding to a *bona fide* FRET response towards lower [MgATP^2-^], as described by De Col *et al*. (2017) for ATeam1.03-nD/nA. In a low pH environment, the proportion of MgATP^2-^ species in a multi-component solution can turn to almost zero, as exemplified by Storer and Cornish-Bowden (1976). Therefore, the ratio response of ATeam1.03YEMK in buffers below pH 7 is likely indicative of waning [MgATP^2-^] rather than pH sensitivity. Both sensors showed typical changes of their emission spectrum in response to increasing [ATP] in accordance to FRET response, as expected (Fig. 2C-D). With increasing [ATP], CFP fluorescence decreased, while YFP fluorescence increased. Within seconds, both sensors showed a rapid response to changing [ATP], and equilibration within 1 minute (Fig. 2E-F).

**Fig. 2:**
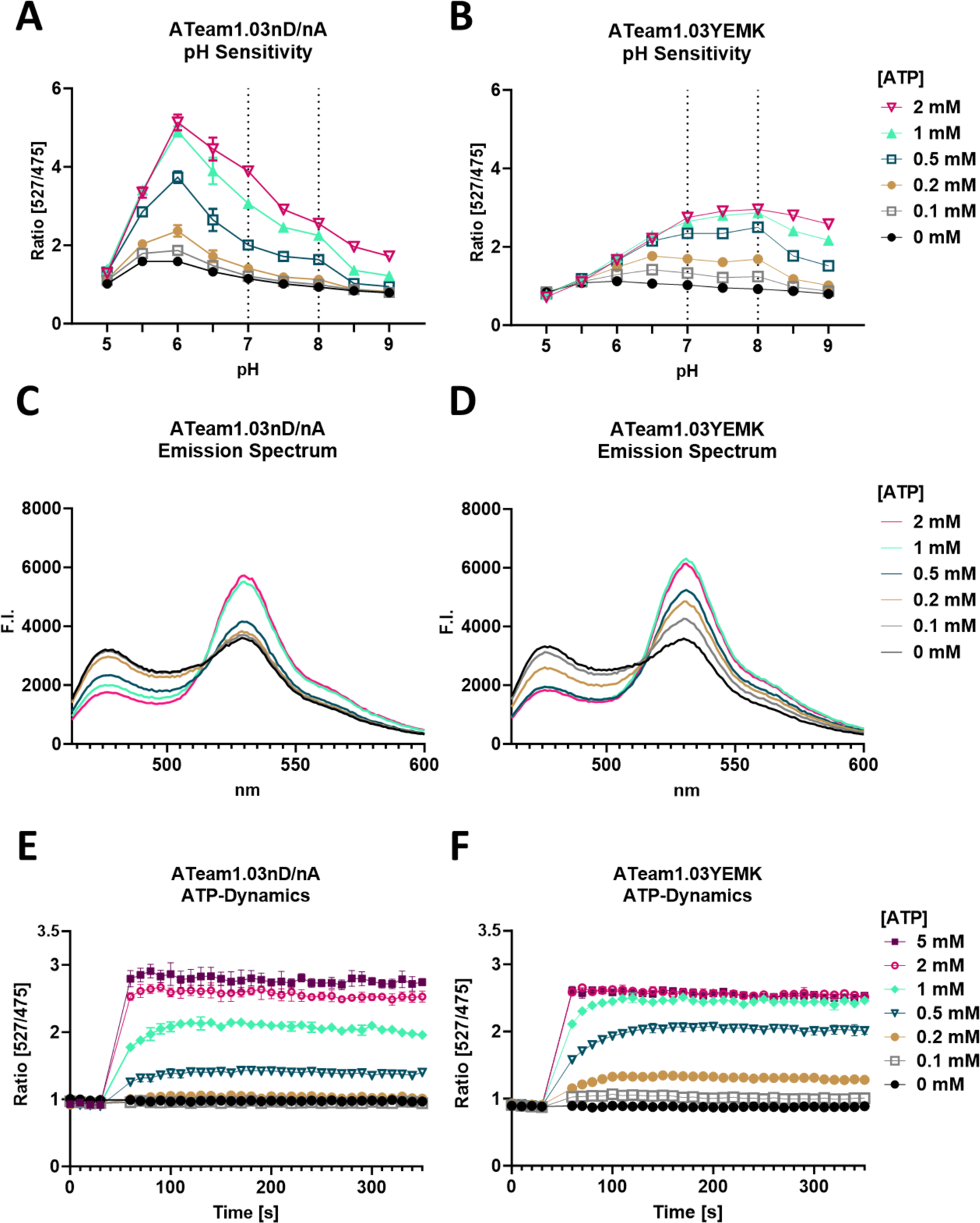
pH, emission spectrum, and time-response characteristics of ATeam1.03-nD/nA and ATeam1.03YEMK. A-B: ATeam1.03-nD/nA shows a higher dynamic range but less pH stability than ATeam1.03YEMK; C-D: Both sensors show a characteristic change of emission spectrum after excitation at 435 and emission detection from 463 - 600 nm in response to different [ATP] in accordance to FRET. Error bars were omitted for clarity; E-F: Both sensors show a rapid response towards changes in [ATP]. The ratio reached a stable level approximately 1 minute after ATP addition; Measurements were conducted via excitation at 435 nm and emission sensing at 527 nm for YFP and 475 nm for CFP. Mean values of n = 3 independent experiments are shown. Error bars indicate SD.

ATeam fluorescence and FRET response is influenced by temperature and pH (Imamura *et al*. 2009). We aimed to imitate the environment in living parasites and therefore chose 37 °C and pH 7.34 (cytosolic pH as determined in later experiments (Fig. 9B)) for all experiments with recombinant protein, if not indicated otherwise. Drug-sensor interaction studies did not find any relevant direct interaction between compounds and sensors in our desired concentration range for ATeam1.03YEMK (Fig. S2A) and ATeam1.03-nD/nA (data not shown).

### ATeams can be stably expressed in *P. falciparum* NF54*attB* and are capable of measuring dynamic real-time changes of ATP level in a plate reader format

We used the NF54*attB* cell line lacking a selectable marker (Adjalley *et al*. 2011) to generate our transgenic cell lines. This allows *attB*-*attP* recombination within the cg6 gene using the mycobacteriophage Bxb1 integrase (Nkrumah *et al*. 2006), and thereby, facilitates rapid generation of stable transfectants. We used limiting dilution to generate clonal lines and used PCR for verification (Fig. S3, Fig. S4). NF54*attB*^[ATeam1.03-nD/nA]^ and NF54*attB*^[ATeam1.03YEMK]^ cell lines showed sensor-specific fluorescence in a shape resembling the cytosol of the parasites, as shown for a representative NF54*attB*^[ATeam1.03-nD/nA]^ clonal line (Fig. 3A-E) with similar absolute fluorescence intensity within each cell population (data not shown).

**Fig. 3:**
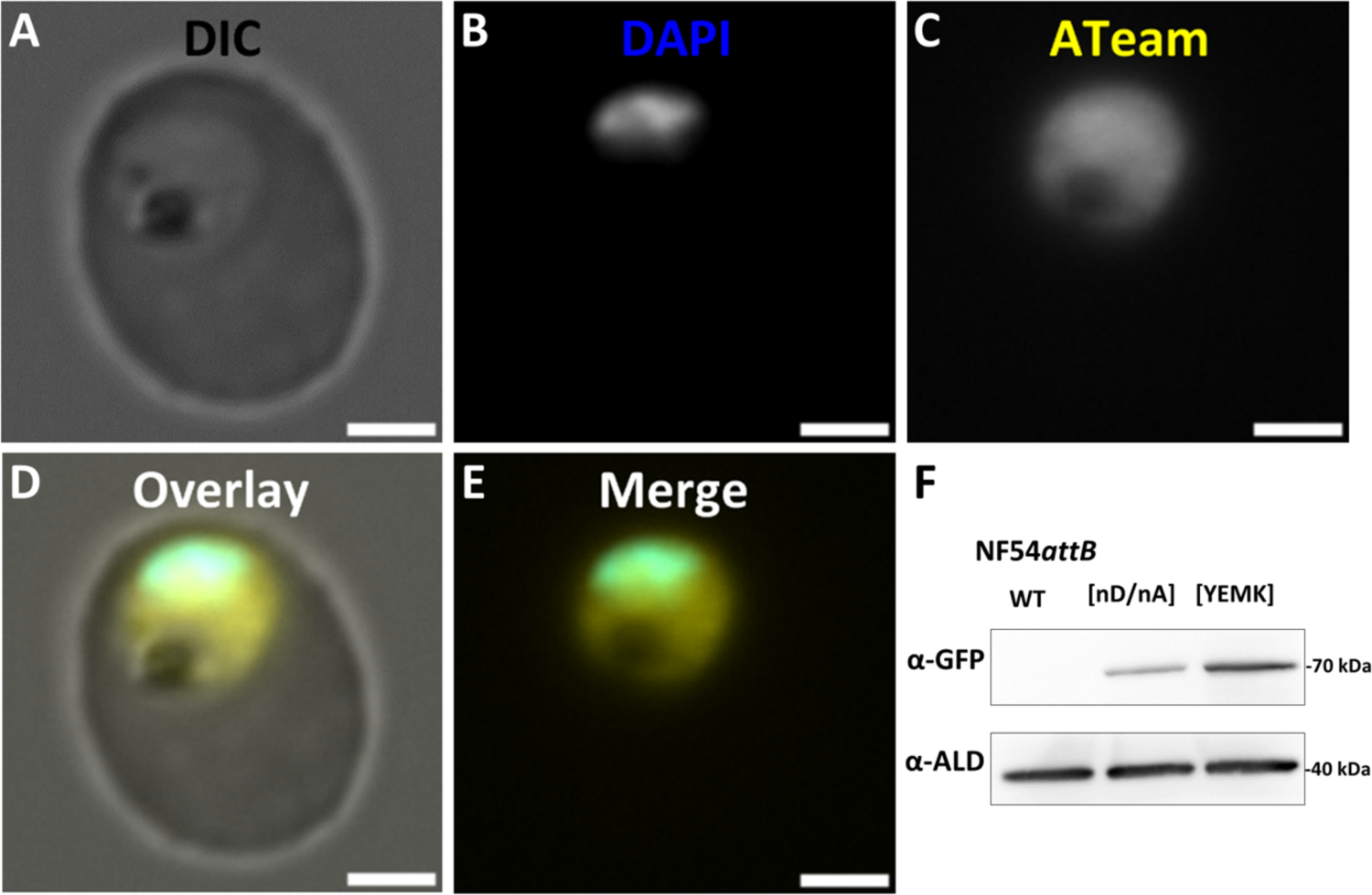
NF54*attB*^[ATeam1.03YEMK]^ and NF54*attB*^[ATeam1.03-nD/nA]^ show cytosolic ATeam fluorescence. DIC (A), DAPI (B), ATeam (C), Overlay of A-C (D) and merge of B+C (E) of NF54*attB*^[ATeam1.03-nD/nA]^ as representative for both ATeam cell lines. DAPI: Ex 353 nm/ Em 465 nm. ATeam: Ex 488 nm / Em 509 nm. Scale bar equals 2 µm; F: Both cell lines show α-GFP signal with the size of ATeam1.03-nD/nA and ATeam1.03YEMK protein (67.8 and 68.0 kDa, respectively). NF54*attB*^[ATeam1.03YEMK]^ shows 1.96 ± 0,38 (SD) fold higher western blot signal than NF54*attB*^[ATeam1.03-nD/nA]^ (n =3) normalized to α-Aldolase as loading control.

Under identical imaging conditions, NF54*attB*^[ATeam1.03YEMK]^ clonal lines showed constantly higher absolute fluorescence intensity than NF54*attB*^[ATeam1.03-nD/nA]^ clonal lines (data not shown). As ATeams are based on CFP and YFP proteins that are derived from GFP, we used cross-reactive anti-GFP antibodies for western blot analyses. In accordance to lower fluorescence intensity, western blot analysis revealed a higher signal intensity of NF54*attB*^[ATeam1.03YEMK]^ compared to NF54*attB*^[ATeam1.03-nD/nA]^ relative to aldolase loading control, indicating higher sensor abundance in the NF54*attB*^[ATeam1.03YEMK]^ parasites (Fig. 3F).

To demonstrate proof-of-principle of ATeam-expressing cell lines in a plate reader format, we analysed their emission spectrum after glucose (GLC) starvation in response to different GLC solutions. As the parasite’s metabolism is heavily reliant on GLC, GLC starvation is expected to cause ATP depletion and an according ATeam emission spectrum, while GLC addition after starvation is expected to restore ATP level, with increasing YFP and decreasing CFP emission peaks. NF54*attB*^[ATeam1.03-nD/nA]^ and NF54*attB*^[ATeam1.03YEMK]^ showed a change in emission spectrum with increasing [GLC] and a 10 minute incubation time, in accordance with the FRET response (Fig. 4A+B). A marked increase in YFP and decrease in CFP peak was already observable with 150 µM GLC. There is no apparent difference between addition of 1 and 20 mM GLC. No conditions fully replicated the emission spectrum of the control spectrum of parasites that were kept continuously in GLC-rich Ringer’s solution (Control). NF54*attB*^[ATeam1.03-nD/nA]^ showed overall lower fluorescence intensity compared to NF54*attB*^[ATeam1.03YEMK]^ under comparable conditions, leading to a lower signal to noise ratio.

**Fig. 4:**
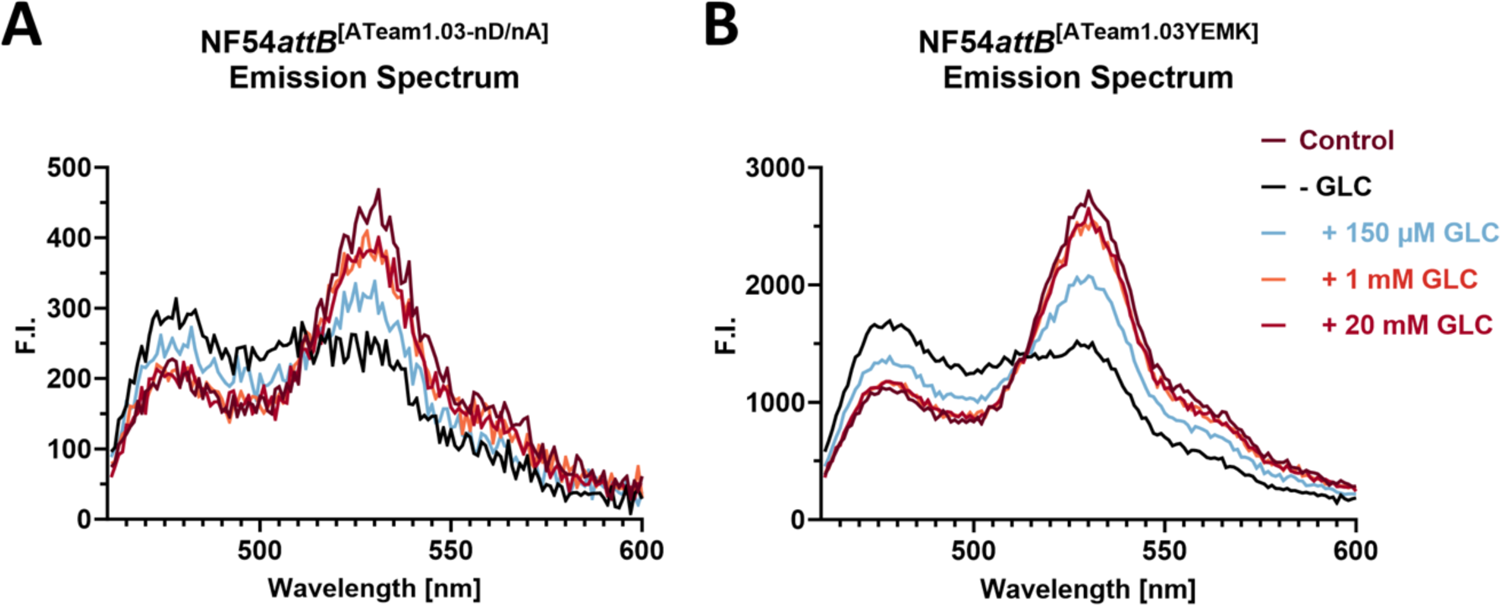
Emission spectrum of NF54*attB*^[ATeam1.03-nD/nA]^ and NF54*attB*^[ATeam1.03YEMK]^ responds to glucose titration in accordance to ATeam FRET. A: NF54*attB*^[ATeam1.03-nD/nA]^; B: NF54*attB*^[ATeam1.03YEMK]^; Plate reader emission spectra of 2 million MACS-enriched trophozoite-infected red blood cells (iRBCs) cells after excitation at 435 nm and emission detection from 463 - 600 nm in response to different [GLC] after 10 minutes incubation time are shown. Parasites were washed in PBS for GLC depletion (-GLC) or GLC-rich Ringer’s solution (Control). Mean emission spectrum of n = 3 independent experiments are shown. Error bars were omitted for clarity.

We further demonstrated proof-of-principle of dynamic plate reader measurements over time using treatment with GLC and subsequent addition of the glycolysis inhibitor 2-desoxyglucose (2-DG [Wick *et al*. 1957]) after GLC starvation (Fig. 5). GLC addition is expected to restore ATP levels, while addition of 2-DG is expected to deplete parasite ATP levels. Cells were washed in PBS for GLC depletion or GLC-rich Ringer’s solution as a control. After 2 minutes, cells were treated with either PBS, Ringer’s, or 1 mM GLC solution. Within 10 minutes, GLC-depleted cells treated with 1 mM GLC solution restored their emission ratio almost to that of the control group in Ringer’s solution, indicating a restoration of their ATP level. The ratio matched that of the control group in Ringer’s solution over an additional 30 minutes within the plate reader, while both groups show a small but steady ratio decline over time within the plate reader. In contrast, addition of 2-DG 10 minutes after recovery of GLC-depleted cells caused a rapid ratio decrease to the baseline level in GLC-depleted PBS solution. Under the same conditions, NF54*attB*^[ATeam1.03-nD/nA]^ showed a similar response but much lower signal to noise ratio (Fig. 5B). As both total sensor fluorescence and pH independence were lower when using ATeam1.03-nD/nA, we decided to prioritize ATeam1.03YEMK for all further experiments.

**Fig. 5:**
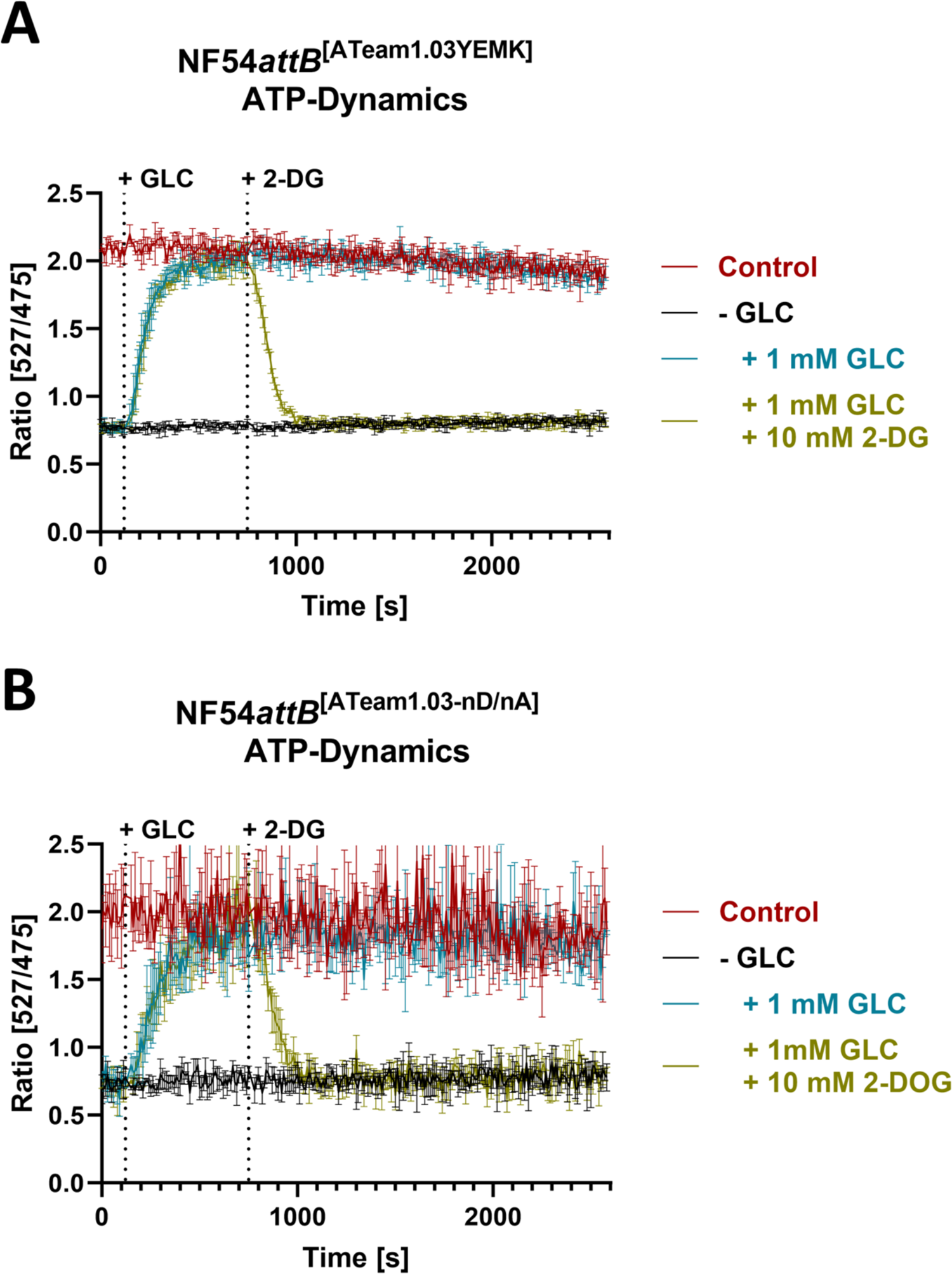
Monitoring of NF54*attB*^[ATeam1.03YEMK]^ and NF54*attB*^[ATeam1.03-nD/nA]^ emission ratio demonstrates dynamic *in cellulo* sensor response in plate reader format. Plate reader emission ratio of 2 million MACS-enriched NF54*attB*^[ATeam1.03YEMK]^ (A) or NF54*attB*^[ATeam1.03-nD/nA]^ (B) trophozoite-iRBCs after excitation at 435 nm and emission sensing at 527 nm for YFP and 475 nm for CFP. Parasites were washed in PBS for GLC depletion (-GLC) or GLC-rich Ringer’s solution (Control). Treatment with 1 mM GLC restored ratio to control in Ringer’s solution. Additional treatment with 10 mM 2-DG after 10 minutes reversed GLC-induced restoration to GLC-deprived baseline ratio. Mean emission ratio of n = 3 independent experiments are shown. Error bars indicate SD.

### NF54a*ttB*^[ATeam1.03YEMK]^ can be used to monitor dynamic changes of ATP at single-cell level using fluorescence microscopy

Following on from plate reader measurements, we established NF54a*ttB*^[ATeam1.03YEMK]^ measurements on a single-cell level using fluorescence microscopy. Similar to the plate reader experiments (Fig. 5), we treated GLC-starved parasites with GLC and subsequent addition of 2-DG. Fig. 6 shows the emission ratio, overlay, YFP- and CFP-emission channel of a representative trophozoite infected red blood cell (iRBC) over time. The cell dynamically reacts to GLC addition with increase and decrease of fluorescence intensities in the YFP and CFP channel, respectively, leading to an increase in emission ratio. The process is reversed via application of 2-DG. Plotting the emission ratio of multiple single cells over time demonstrates a dynamic *in cellulo* sensor reaction in a fluorescence microscope (Fig. 7). The measurements also reveal a steady emission ratio decrease of untreated control parasites that may reflect limited photo bleaching.

**Fig. 6:**
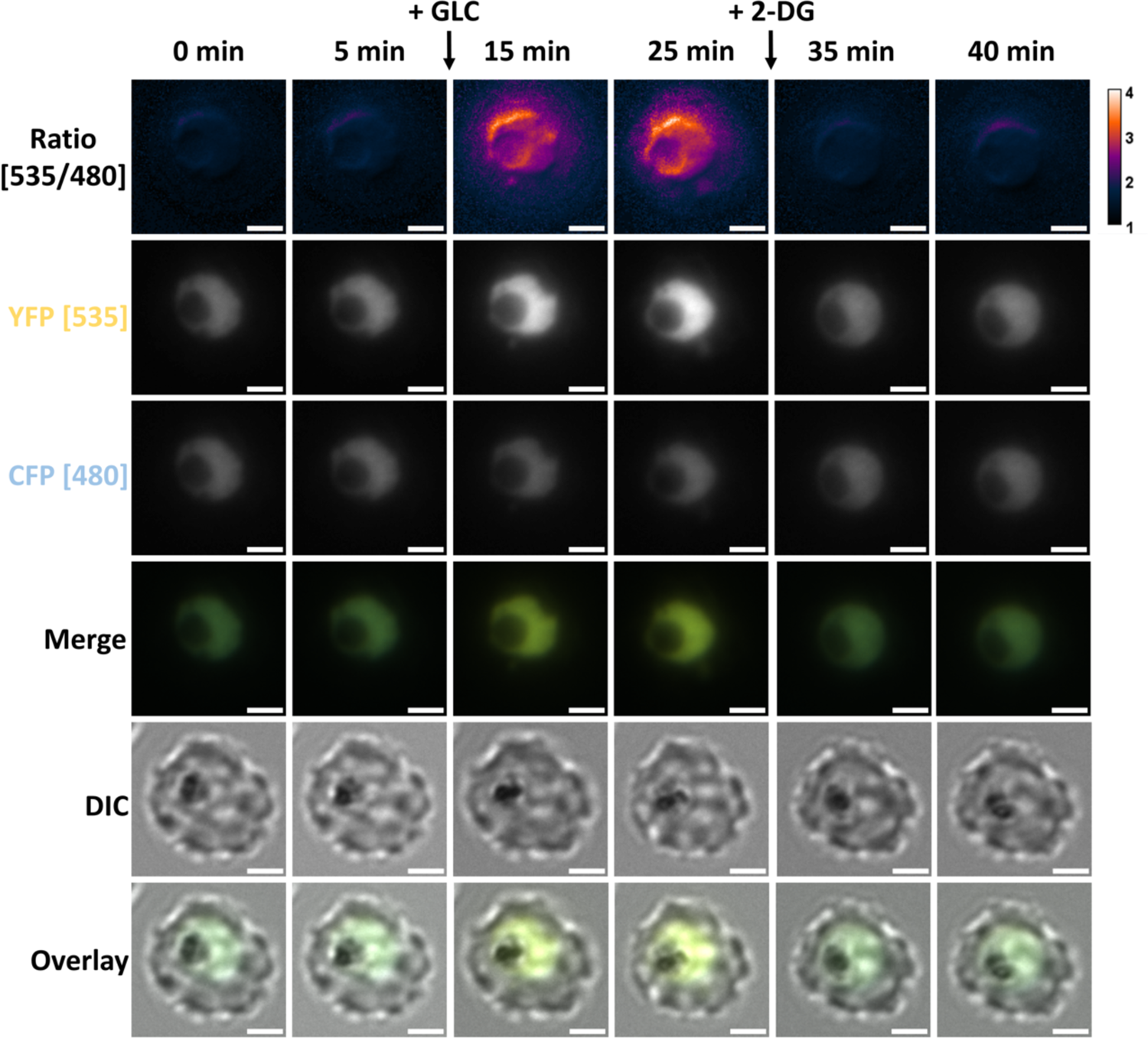
Fluorescence microscopy of NF54a*ttB*^[ATeam1.03YEMK]^ allows monitoring of ATP-glucose dynamics on single-cell level. Emission ratio of NF54*attB*^[ATeam1.03YEMK]^ was measured using excitation at 430 nm and emission sensing at 535 nm and 480 nm for the YFP and CFP channel, respectively. GLC was added to a concentration of 2 mM after taking the 5 min image. 2-DG was added to a concentration of 10 mM after taking the 25 min image. Scale bar equals 2 µm.

**Fig. 7:**
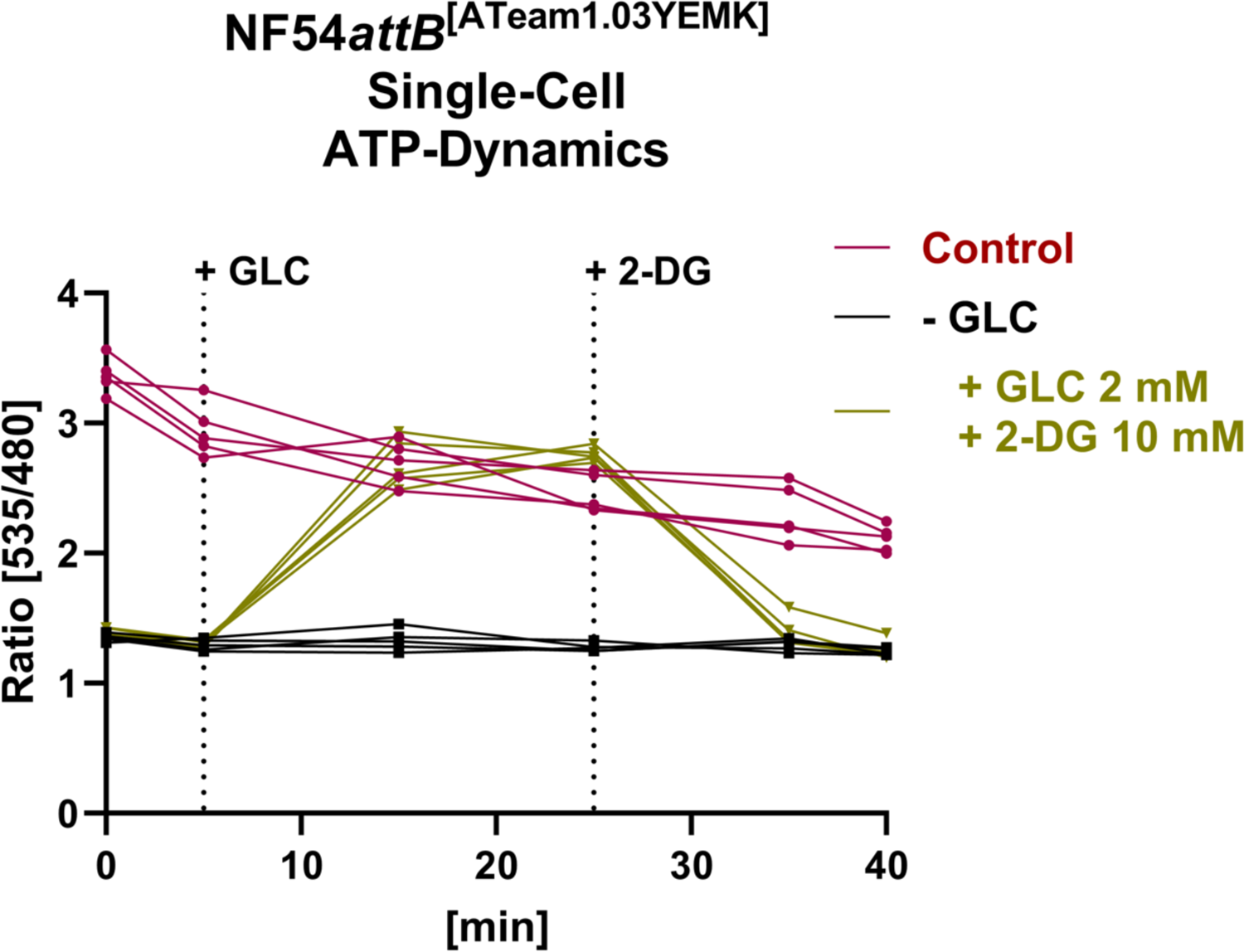
Fluorescence microscopy of NF54a*ttB*^[ATeam1.03YEMK]^ demonstrates dynamic *in cellulo* sensor response on single-cell level. Emission ratio of NF54*attB*^[ATeam1.03YEMK]^ was measured using excitation at 430 nm and emission sensing at 535 nm and 480 nm for the YFP and CFP channel, respectively. Background-corrected ratios are shown. Cells were washed and treated either with GLC-rich Ringer’s solution (Control) or GLC-depleted PBS (-GLC). Intervention group was treated with GLC and 2-DG after GLC starvation in PBS as indicated. GLC was added to a concentration of 2 mM after taking the 5 min image. 2-DG was added to a concentration of 10 mM after taking the 25 min image. Continuous measurements of n = 5 parasites/group is shown.

### The pH sensor sfpHluorin allows robust measurement of cytosolic pH and is capable of measuring dynamic pH changes in a plate reader

As shown in Fig. 2, *in cellulo* ATeam measurements could be affected by pH fluctuations. To control for this, we generated parasites expressing the genetically-encoded pH sensor sfpHluorin. Additionally, we produced His-tagged sfpHluorin protein to characterize the sensor’s behavior *in vitro* and to control for possible direct drug-sensor interactions. pH titration and changes of emission and excitation spectra in a plate reader (Fig. S5) was in accordance with Reifenrath and Boles (2018). We could not detect any direct drug-sensor interactions in our desired concentration range (Fig. S2B). We generated stable sfpHluorin-expressing cell lines and used limiting dilution to isolate clonal lines. To verify stable integration we used PCR (Fig. S6). The NF54*attB*^[sfpHluorin]^ clonal line showed sensor-specific fluorescence in a shape resembling the parasitic cytosol (Fig. 8).

**Fig. 8:**
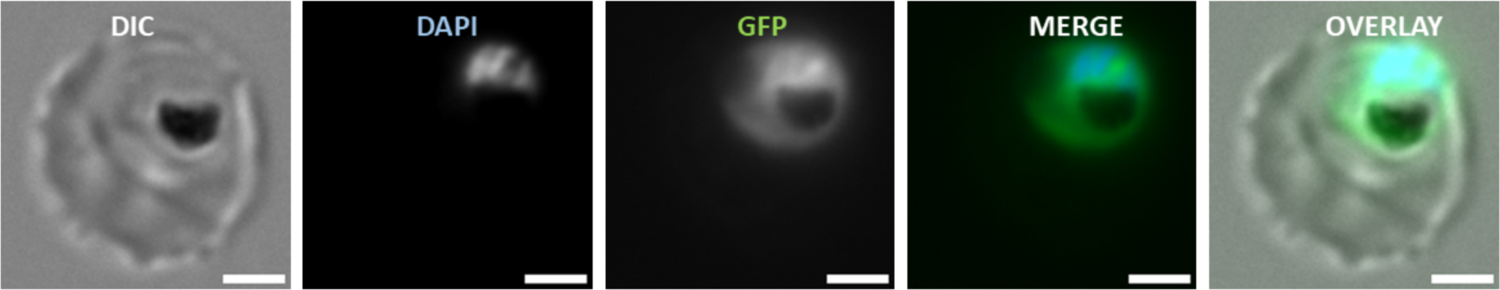
NF54*attB*^[sfpHluorin]^ shows cytosolic GFP fluorescence. DIC, DAPI, GFP, merge, and overlay of a representative trophozoite stage parasite of NF54*attB*^[sfpHluorin]^ is shown. DAPI: Ex 353 nm/ Em 465 nm. GFP: Ex 488 nm / Em 509 nm. Error bar equals 2 µm.

To demonstrate proof-of-principle of NF54*attB*^[sfpHluorin]^ *in cellulo*, we applied nigericin pH calibration. Nigericin is an ionophore, allowing pH equilibration of the cytosol with the extracellular media (Thomas *et al*. 1979). Spectral analysis of nigericin-permeabilized NF54*attB*^[sfpHluorin]^-iRBCs reveals excitation spectra changing in response to pH in accordance with that of recombinant sfpHluorin, with pH-responding peaks around 390 and 482 nm (Fig. 9A). The excitation ratio of 390 to 482 nm plotted over pH level allows fitting of a sigmoidal calibration curve for *in cellulo* pH measurements with a high dynamic range in the physiological relevant range of the parasitic cytosol between pH 6 and 8. Estimation of parasite resting pH in physiologic Ringer’s solution determines a pH ± SD of 7.34 ± 0.07 (Fig. 9B), in large agreement with previous determinations, such as 7.29 ± 0.01 (Saliba and Kirk 1999), 7.31 ± 0.02 (Hayashi *et al*. 2000), or 7.3 ± 0.05 (Wissing *et al*. 2002).

**Fig. 9:**
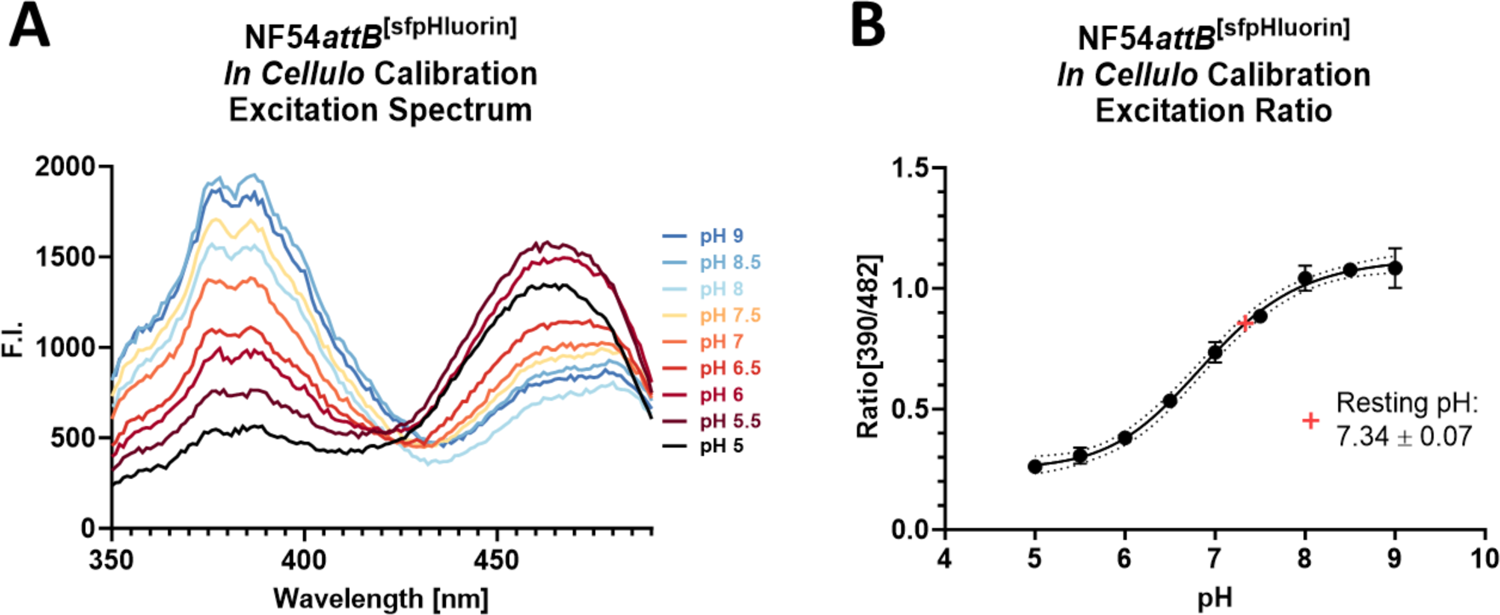
NF54*attB*^[sfpHluorin]^ *in cellulo* calibration in plate reader format. 1 million MACS-enriched trophozoite-iRBCs incubated with 10 µM nigericin for 30 minutes at 37 °C in 384-well plates in buffers with pH from 5 to 9. A: Excitation spectra from 350 to 490 nm with emission sensing at 530 nm show pH responsive peaks around 390 and 482 nm. Error bars were omitted for clarity; B: Fitting a calibration curve using excitation ratio of 390 to 482 nm with emission sensing at 530 nm of nigericin-treated cells in pH buffers allows pH determination of untreated cells in Ringer’s solution. Means of n = 3 independent experiments is shown. Error bars indicate SD.

To further demonstrate the suitability of NF54*attB*^[sfpHluorin]^ for measuring dynamic pH changes, the cell line underwent GLC starvation, recovery, and subsequent 2-DG treatment as in previous experiments (Fig. 10). Starvation from GLC in PBS led to a lower excitation ratio compared to the control. Treatment with 1 mM GLC restored the excitation ratio to almost that of the control. Subsequent 2-DG treatment after 10 minutes rapidly reversed the GLC-induced recovery. The dynamic pH response of the parasite in response to GLC starvation and recovery shown here matches the findings of Saliba and Kirk (1999) who showed that starvation from GLC causes a drop in parasite cytosolic pH, and that recovery from acidification is heavily impaired under GLC starvation.

**Fig. 10:**
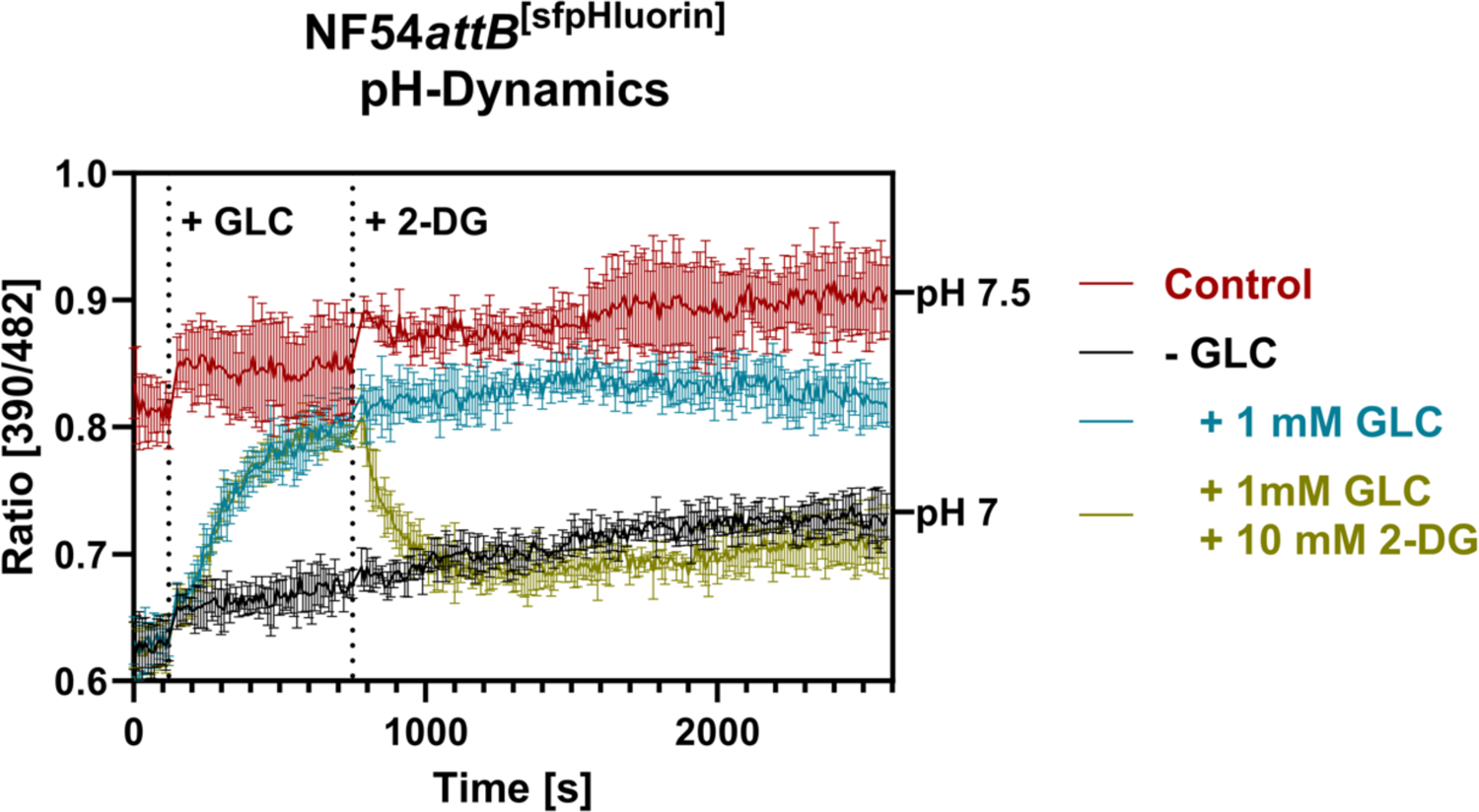
Monitoring of NF54*attB*^[sfpHluorin]^ excitation ratio demonstrates dynamic *in cellulo* sensor response in plate reader format. Plate reader excitation ratio of 1 million MACS-enriched trophozoite infected-iRBCs after excitation at 390 to 482 nm with emission sensing at 530 nm. Parasites were washed in PBS for GLC depletion (-GLC) or GLC-rich Ringer’s solution (Control). Treatment with 1 mM GLC restored ratio close to control in Ringer’s solution. Additional treatment with 10 mM 2-DG after 10 minutes reversed GLC-induced restoration close to GLC-starved baseline ratio. Mean emission ratio of n = 3 independent experiments are shown. Error bars indicate SD.

### NF54*attB*^[sfpHluorin]^ can be used to monitor pH changes at a single-cell level using live-cell imaging

To control for pH changes that might occur during sample preparation and measurements in the single-cell microscopy setup for NF54*attB*^[ATeam1.03YEMK]^, we further established NF54*attB*^[sfpHluorin]^ measurements in a similar live-cell setup. We used the nigericin calibration method as described earlier. Accordingly, we observed a 385 to 475 nm excitation ratio increase with increasing pH and a high dynamic range between pH 6 and 8. Through interpolation of a standard curve, we could determine the resting pH (mean ± SD) of parasites in the microscopy setup in Ringer’s solution to be 7.32 ± 0.12 (Fig. 11A), in strong accordance to our measurements made using a plate reader (7.34 ± 0.07, Fig. 9B).

**Fig. 11:**
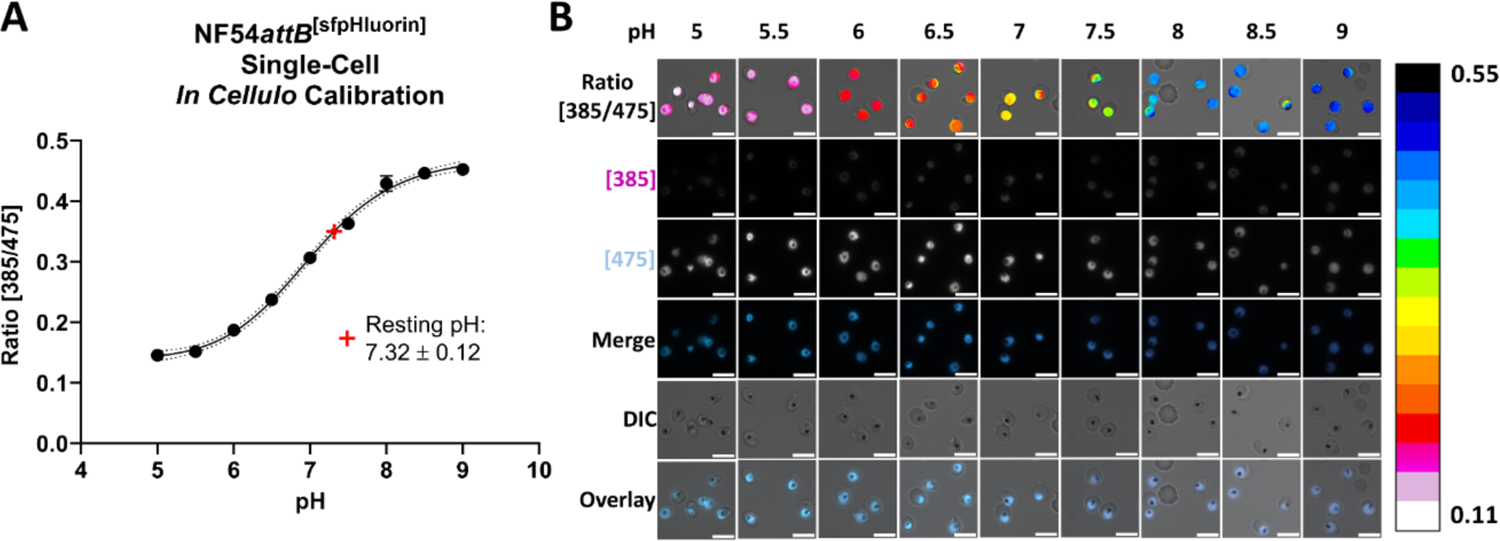
NF54*attB*^[sfpHluorin]^ single-cell *in cellulo* calibration in epifluorescence microscope. MACS-enriched trophozoite-iRBCs incubated with 10 µM nigericin for 30 minutes at 37 °C in buffers with pH from 5 to 9 show pH responsive increase in 385 to 475 nm excitation ratio at 525 nm. A: Fitting a calibration curve allows resting pH determination of untreated cells in Ringer’s solution. Measurements show mean of n = 3 independent experimental means of 100 cells ± SD; B: Representative single-cell images of calibration experiments. Calibration bar color codes pixel by pixel 385 to 475 nm excitation ratio at 525 nm. Scale bar equals 10 µm.

### NF54*attB*^[ATeam1.03YEMK]^ shows distinct response patterns towards different drug classes

Having established NF54*attB*^[ATeam1.03YEMK]^ and NF54*attB*^[sfpHluorin]^ as ATP and pH measurement systems, we wished to use these new tools to study parasite responses to different established drug classes and promising drug candidates. We incubated the sensor cell lines with the compounds at approximately 100x and 10x of their EC_50_ concentrations (Table S1) to induce rapidly detectable effects over an incubation time of 4 h and 6 h. Exceptions to this approach were doxycycline (DOXY) and the model inhibitor of protein synthesis cycloheximide (CHX). DOXY was used at the fixed concentrations 5 µM and 1 µM to account for first and second cycle effects (Dahl *et al*. 2006), and CHX was used at the single dose of 50 µg/mL to demonstrate the effect of a protein synthesis inhibitor.

To detect the ATeam response, we used the microscopic measurement approach with n = 5 independent experiments and calculated the mean of 100 parasites for each independent experiment with a concurrent vehicle-treated control group in each microscopic dish. We used this randomized block design to tease out parasite responses to different compound classes and used two-way ANOVA for statistical analysis. To cope with the multiple comparisons problem, we used the false discovery rate (FDR [Benjamini *et al*. 2006]). Discoveries with 5 % FDR are reported with a single asterisk.

After 6 h incubation, we found distinct ATeam response towards different drug classes (Fig. 12). The 4-aminoquinoline compounds chloroquine (CQ), amodiaquine (AQ), and pyronaridine (PYRO) caused a marked drop in ATeam emission ratio at 100x of their EC_50_ and, except for CQ, additionally at 10x EC_50_. In contrast, the arylamino alcohols mefloquine (MQ) and lumefantrine (LUM) caused an increase in emission ratio, both at 100x and 10x EC_50_ concentration. The structurally related quinine (QN) did not show any effect on ATeam emission ratio. Dihydroartemisinin (DHA) caused a mixed effect on the emission ratio. We could detect a decrease in emission ratio for 100x EC_50_, while we could detect an increase for 10x EC_50_. The redox-active compounds methylene blue (MB [Schirmer *et al*. 2011]) and plasmodione (PD [Ehrhardt *et al*. 2016]) caused an emission ratio decrease at 100x EC_50_, but not at 10x EC_50_. The apicoplast effecting drug (Dahl *et al*. 2006) DOXY, the inhibitor of the mitochondrial ETC (Fry and Pudney 1992) ATQ, as well as the PfGluPho inhibitor SBI-0797750 (Berneburg *et al*. 2022) did not show any effects under the given conditions. Additionally, we included the effects of GLC starvation (-GLC) and the inhibitor of protein biosynthesis CHX in our study. As expected, GLC starvation just before measurement caused a drastic drop in emission ratio. In contrast, CHX caused an increase in ratio. Overall, we found similar results for all tested compounds already after 4 h incubation. However, the results were less pronounced and not detected as discovery according to the FDR for 10x EC_50_ of AQ, PYRO, MQ, DHA, SBI, and 100x and 10x EC_50_ of LUM (Fig. S7).

**Fig. 12:**
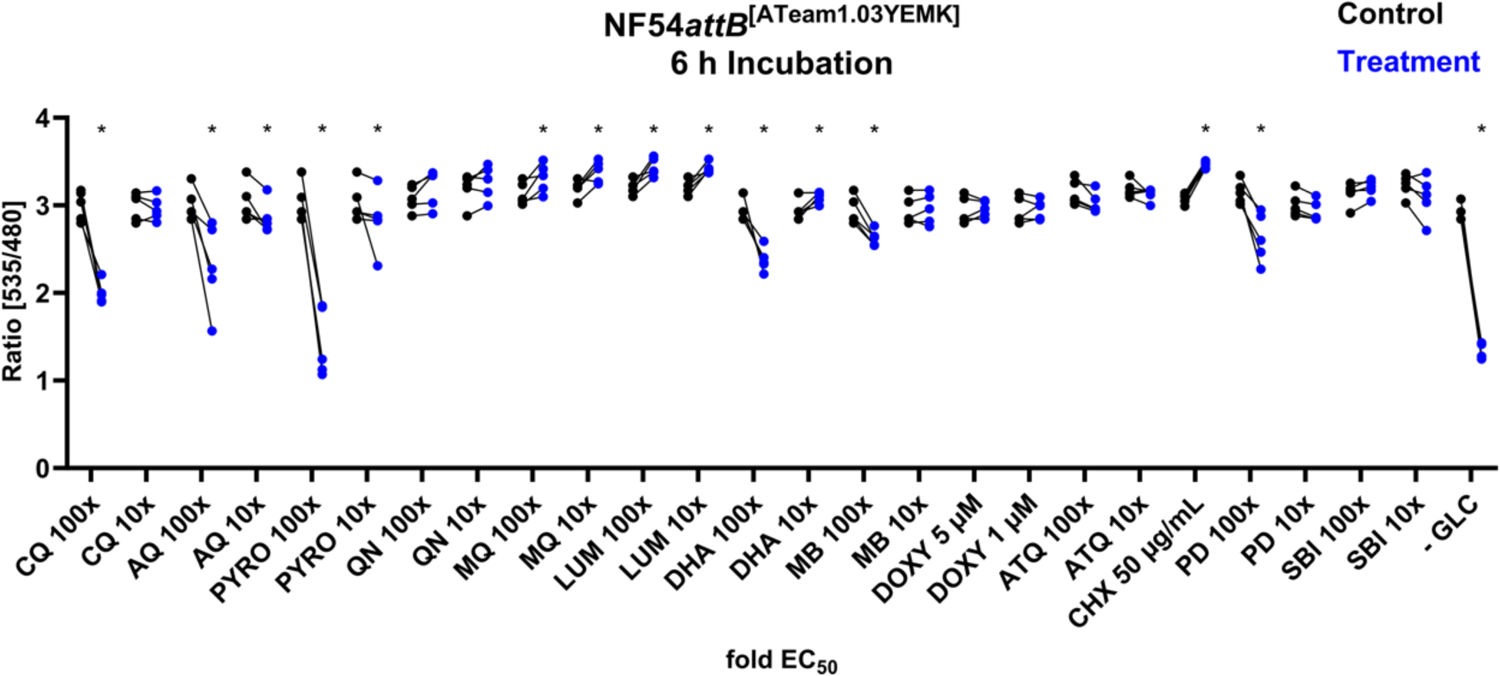
NF54*attB*^[ATeam1.03YEMK]^ shows distinct ratio changes in response towards different drug classes after 6 h incubation. Fluorescence microscopic measurement of MACS-enriched trophozite-iRBCs. Each measurement corresponds to the 535 nm to 480 nm emission ratio after excitation at 430 nm. Mean of 100 single cell analyses of compound intervention (Treatment) and concurrent vehicle control (Control) with n = 5 independent experiments are shown. Treatment groups included chloroquine (CQ), amodiaquine (AQ), pyronaridine (PYRO), quinine (QN), mefloquine (MQ), lumefantrine (LUM), dihydroartemisinin (DHA), methylene blue (MB), doxycycline (DOXY), atovaquone (ATQ), cycloheximide (CHX), plasmodione (PD), SBI 0797750 (SBI), and glucose starvation (-GLC). Cells were starved from GLC through washing in PBS just before measurement. If not indicated otherwise, compounds were applied with 100x and 10x EC_50_ concentration. * indicates discovery with FDR = 5 %.

Pooling the single-cell data of the n = 5 independent experiments gives further insights into the compound-induced ATeam response. Though not suitable for statistical analysis, as the pooling and lack of concurrent control would make statistical analysis questionable, we nevertheless observed distinct ATeam response patterns towards different drug classes on a single-cell level (Fig. 13). The 4-aminoquinolines CQ, AQ, and PYRO, caused a distinctive bimodal ratio distribution at 100x EC_50_ concentration. A notable proportion of their ratio distributions are located below 1, while even cells starved of GLC did not show such low ratios. In contrast, the distributions of the arylamino alcohols QN, MQ, LUM, in addition to CHX, and SBI are more compressed and higher located compared to a representative control group. DHA caused mixed effects in the ratio distribution. It showed a compressed and higher located distribution at 10x EC_50_, while it shows tendency towards a bimodal and downwardly skewed distribution at 100x EC_50_. MB showed a tendency for compression and lower ratios at 100x EC_50_. PD showed tendency towards lower and bimodal distribution. DOXY and ATQ did not show any apparent change of their distributions compared to the representative control.

**Fig. 13:**
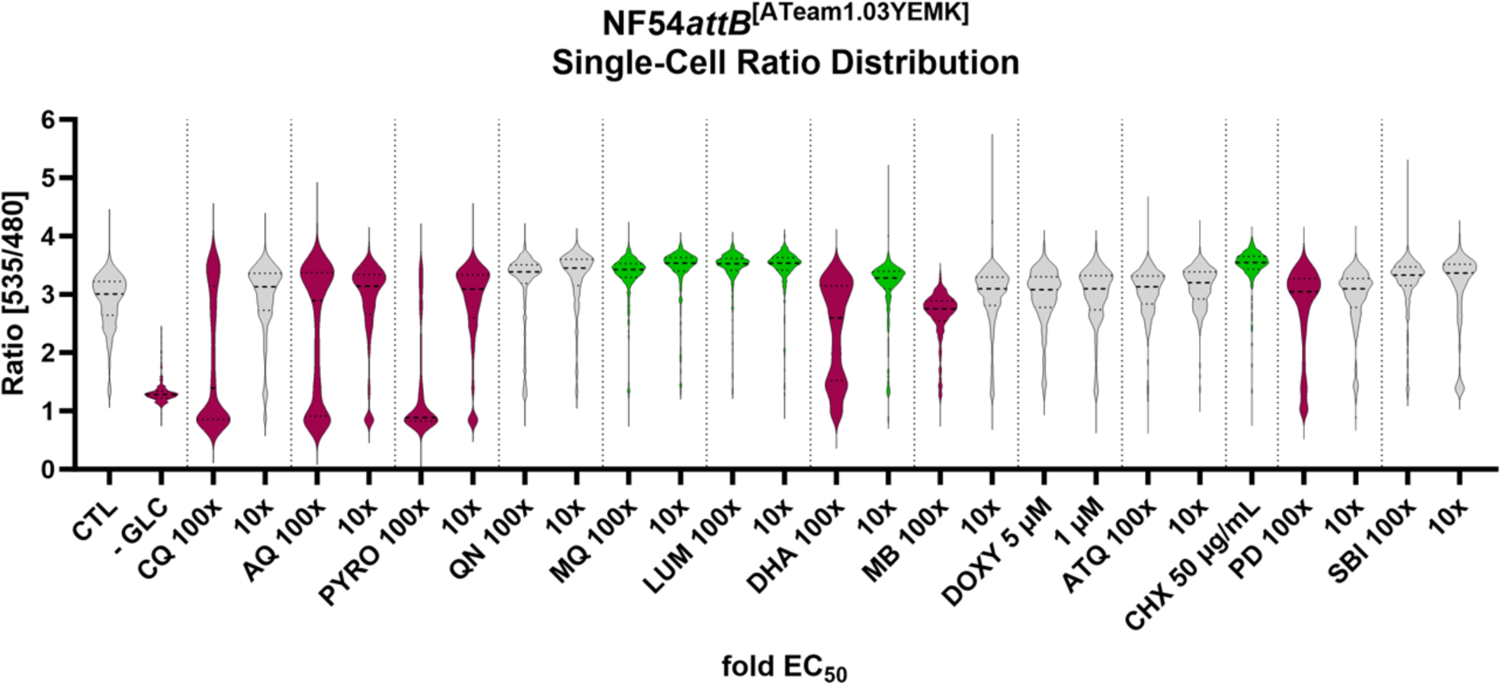
NF54*attB*^[ATeam1.03YEMK]^ shows distinct single-cell emission ratio distribution in response towards different drug classes. Fluorescence microscopic measurement of MACS-enriched trophozite-iRBCs. Each measurement corresponds to the 535 nm to 480 nm emission ratio after excitation at 430 nm. Pool of n = 5 independent experiments with each 100 single cell analyses is shown. Treatment groups included chloroquine (CQ), amodiaquine (AQ), pyronaridine (PYRO), quinine (QN), mefloquine (MQ), lumefantrine (LUM), dihydroartemisinin (DHA), methylene blue (MB), doxycycline (DOXY), atovaquone (ATQ), cycloheximide (CHX), plasmodione (PD), SBI 0797750 (SBI), and glucose starvation (-GLC). Cells were starved from GLC through washing in PBS just before measurement. If not indicated otherwise, compounds were applied with 100x and 10x EC_50_ concentration. Colors indicate up (green) or down (magenta) shifted discoveries according to statistical analysis described in Fig. 12.

### pH changes induced by drug interventions are generally small and unlikely to affect *in cellulo* ATeam measurements

As shown in Fig. 2B, ATeam1.03YEMK showed a stable ratio response within the pH range 6-8. In order to investigate if pH changes might influence ATeam measurements upon treatment, all compound treatments shown in Fig. 12 were repeated with NF54*attB*^[sfpHluorin]^. We used the microscopic measurement approach with n = 3 independent experiments and calculated the mean of 100 parasites for each independent experiment with a concurrent vehicle-treated control group in each microscopic dish. The pH levels were interpolated from excitation ratio measurements using the *in cellulo* calibration shown in Fig. 11. After 6 h incubation, we found distinct sfpHluorin responses towards different drug classes (Fig. 14). The 4-aminoquinoline compounds CQ, AQ, PYRO, as well as PD at their 100x EC_50_ concentration caused a marked drop below pH 7. Among all compounds tested, only DOXY at a concentration of 5 µM caused an increased pH, however this is still in the range classed as reliable for ATeam measurements. All other compounds caused no or only marginal sfpHluroin excitation ratio changes through the concentrations tested. In addition to that, GLC-starvation just before measurement caused the strongest excitation ratio drop, indicating a mean pH ± SD of 6.71 ± 0.02. With the exception of CQ and LUM 10x EC_50_ concentrations, we could detect the same significant trends after only 4 h incubation time (Fig. S8).

**Fig. 14:**
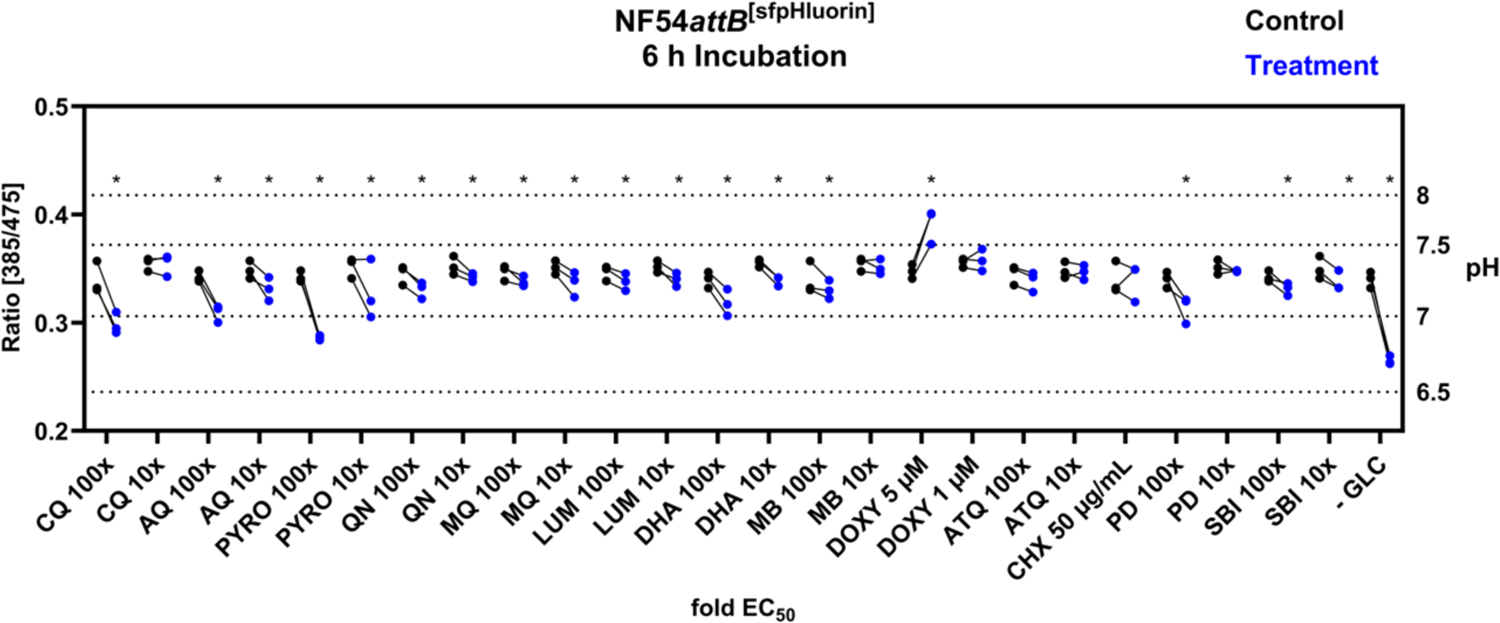
NF54*attB*^[sfpHluorin]^ shows predominantly marginal ratio changes in response towards different drug classes after 6 h incubation time. Fluorescence microscopic measurement of MACS-enriched trophozite-iRBCs. Each measurement corresponds to the 385 to 475 nm excitation ratio at 525 nm emission. Mean of 100 single cell analyses of compound intervention (Treatment) and concurrent vehicle control (Control) with n = 3 independent experiments are shown. Treatment groups included chloroquine (CQ), amodiaquine (AQ), pyronaridine (PYRO), quinine (QN), mefloquine (MQ), lumefantrine (LUM), dihydroartemisinin (DHA), methylene blue (MB), doxycycline (DOXY), atovaquone (ATQ), cycloheximide (CHX), plasmodione (PD), SBI 0797750 (SBI), and glucose starvation (-GLC). Cells were starved from GLC through washing in PBS just before measurement. If not indicated otherwise, compounds were applied with 100x and 10x EC_50_ concentration. * indicates discovery with FDR = 5 %.

As previously demonstrated, ATeam1.03YEMK protein shows a stable ratio response between pH 6 and 8, indicating pH stability of the fluorophores in that range. Although all measured average values lie within this range, plotting the pooled ratio distribution of n = 3 independent experiments with each containing 100 single-cell analyses to the interpolated pH, reveals that some proportion of cells exhibit drastic changes of pH levels, which is masked when analyzing only mean ratios (Fig. S9). For example, at the higher DOXY concentration (5 µM), a number of single-cell measurements lie outside of the calibration range.

### Various compounds cause reduced parasite size

The single-cell image analysis for the NF54*attB*^[ATeam1.03YEMK]^ and NF54*attB*^[sfpHluorin]^ drug response measurements included the definition of ROI that correspond to the area of the parasites. To gain a deeper understanding of the parasite’s response to the compound interventions, we analyzed the mean sizes of these ROI as a proxy for parasite size and compared them to the mean of the control group (Fig. 16). We found that the 4-aminoquinoline compounds CQ, AQ, and PYRO caused a decreased parasite size at their 100x EC_50_ concentrations. Additionally, PYRO caused decreased size already at 10x EC_50_. The arylamino alcohols QN, MQ, LUM, in addition to DHA, and SBI caused a decreased cytosol size at both 100x and 10x EC_50_. In addition, 100x EC_50_ of PD and 50 µg/mL CHX caused reduced size. MB, DOXY and ATQ did not show an effect on parasite size with any of the concentrations tested. Most of the effects were already detected after 4 h incubation. Only AQ 10x, LUM 100x, LUM 10x, and SBI 100x EC_50_ did not show a detectable effect after this shorter incubation time (Fig. S10).

**Fig. 16.**
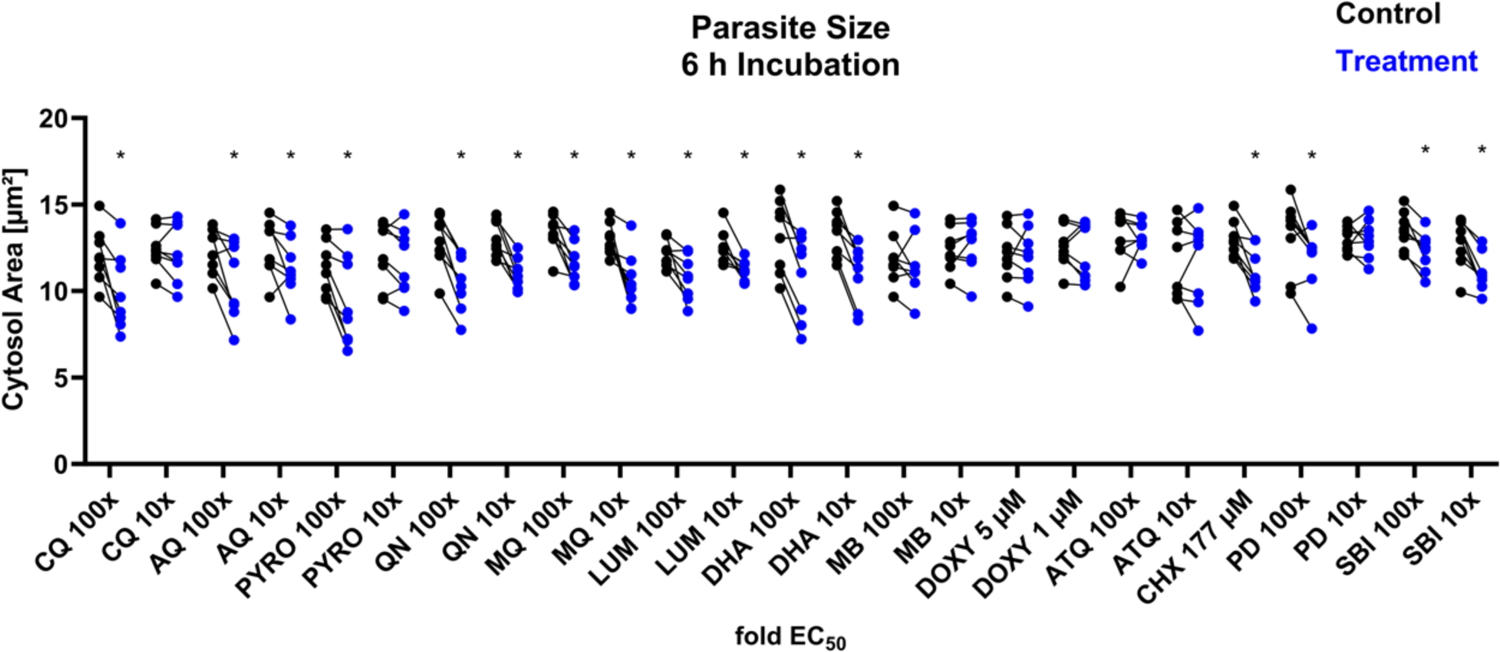
NF54*attB*^[ATeam1.03YEMK]^ and NF54*attB*^[sfpHluorin]^ show decreased parasite size in response towards different drug classes after 6 h incubation. The mean cytosol size of 100 MACS-enriched trophozite-iRBCs derived from automated ROI definition from fluorescence microscopic measurements of NF54*attB*^[ATeam1.03YEMK]^ (n = 5) and NF54*attB*^[sfpHluorin]^ (n = 3) is combined. Analyses of compound intervention (Treatment) and concurrent vehicle control (Control) with in total n = 8 independent experiments are shown. Treatment groups included chloroquine (CQ), amodiaquine (AQ), pyronaridine (PYRO), quinine (QN), mefloquine (MQ), lumefantrine (LUM), dihydroartemisinin (DHA), methylene blue (MB), doxycycline (DOXY), atovaquone (ATQ), cycloheximide (CHX), plasmodione (PD), and SBI 0797750 (SBI). If not indicated otherwise, compounds were applied with 100x and 10x EC_50_ concentration. * indicates discovery with FDR = 5 %.

### Parallel analysis of drug responses reveals common patterns

The characteristics we analysed are not truly independent, as severe metabolic alterations that effect ATP or pH levels are likely to also cause growth effects, and *vice versa.* Accordingly, examination of cytosol size and sensor response patterns in synopsis shows that most of the compounds tested caused either an effect on none, or all of the analysed characteristics after 6 h incubation time (Table 1). CQ 10x, MB 10x, ATQ 100x and 10x, and PD 10x EC_50_, as well as DOXY 1 µM did not show any effects classified as discovery. CQ 100x, AQ 100x and 10x, PYRO 100x, MQ 100x and 10x, LUM 100x and 10x, DHA 100x and 10x, and PD 100x caused effects on all characteristics. PYRO 10x and MB 10x caused effects only on ATeam ratio and pH, while QN 100x and 10x and SBI 100x and 10x only caused effects only on size and pH. CHX only caused effects on cytosol size and ATeam ratio, while DOXY 5 µM caused only an increased pH. Measurements after 4 h incubation showed similar trends, though for some characteristics less pronounced and not detected as discovery.

**Table 1:** Heatmap of sensor and parasite size drug-response patterns shows coinciding effects of compound treatments. The table shows a synopsis of the effects of different compound treatments on size, ATeam emission ratio, and sfpHluorin excitation ratio of NF54*attB*^[ATeam1.03YEMK]^ and NF54*attB*^[sfpHluorin]^ with concentrations in multiples of EC_50_ or otherwise indicated concentrations after 4 or 6 h treatment time. Effects are classified as discovery (Yes), or no discovery (No) with an FDR of 5 %. Color indicates if treatment caused increase (green) or decrease (magenta) of group means, in case a treatment was classified as discovery in analyses described in Fig. 12, 14, 16, S9, S10, S11. Treatment groups included chloroquine (CQ), amodiaquine (AQ), pyronaridine (PYRO), quinine (QN), mefloquine (MQ), lumefantrine (LUM), dihydroartemisinin (DHA), methylene blue (MB), doxycycline (DOXY), atovaquone (ATQ), cycloheximide (CHX), plasmodione (PD), and SBI 0797750 (SBI).

| Discovery? | FDR = 5 % |  |  | Discovery? | FDR = 5 % |  |  |
| --- | --- | --- | --- | --- | --- | --- | --- |
| 6 h | SIZE | FRET | sfpH | 4 h | SIZE | FRET | sfpH |
| CQ 100x | Yes | Yes | Yes | CQ 100x | Yes | Yes | Yes |
| CQ 10x | No | No | No | CQ 10x | No | No | Yes |
| AQ 100x | Yes | Yes | Yes | AQ 100x | Yes | Yes | Yes |
| AQ 10x | Yes | Yes | Yes | AQ 10x | No | No | Yes |
| PYR 100x | Yes | Yes | Yes | PYR 100x | Yes | Yes | Yes |
| PYR 10x | No | Yes | Yes | PYR 10x | No | No | Yes |
| QN 100x | Yes | No | Yes | QN 100x | Yes | No | Yes |
| QN 10x | Yes | No | Yes | QN 10x | Yes | No | Yes |
| MQ 100x | Yes | Yes | Yes | MQ 100x | Yes | Yes | Yes |
| MQ 10x | Yes | Yes | Yes | MQ 10x | Yes | No | Yes |
| LUM 100x | Yes | Yes | Yes | LUM 100x | No | No | Yes |
| LUM 10x | Yes | Yes | Yes | LUM 10x | No | No | No |
| DHA 100x | Yes | Yes | Yes | DHA 100x | Yes | Yes | Yes |
| DHA 10x | Yes | Yes | Yes | DHA 10x | Yes | No | Yes |
| MB 100x | No | Yes | Yes | MB 100x | No | Yes | Yes |
| MB 10x | No | No | No | MB 10x | No | No | Yes |
| DOXY 5 μM | No | No | Yes | DOXY 5 μM | No | No | Yes |
| DOXY 1 μM | No | No | No | DOXY 1 μM | No | No | No |
| ATQ 100x | No | No | No | ATQ 100x | No | No | No |
| ATQ 10x | No | No | No | ATQ 10x | No | No | No |

| CHX 50<br>μg/mL | Yes | Yes | No | CHX 50<br>μg/mL | Yes | Yes | No |
| --- | --- | --- | --- | --- | --- | --- | --- |
| PD 100x | Yes | Yes | Yes | PD 100x | Yes | Yes | Yes |
| PD 10x | No | No | No | PD 10x | No | No | No |
| SBI 100x | Yes | No | Yes | SBI 100x | No | No | Yes |
| SBI 10x | Yes | No | Yes | SBI 10x | Yes | Yes | Yes |
| PBS | NA | Yes | Yes | PBS | NA | Yes | Yes |

### Mefloquine but not amodiaquine prevents chloroquine mediated ATeam and sfpHluorin ratio depletion

We could detect a characteristic drop in ATeam ratio and pH (Table 1) caused by the 4-aminoquinolines CQ, AQ, and PYRO after incubation with 100x EC_50_ for 6 h. CQ, AQ, and PYRO are thought to kill the parasites through inhibition of hemozoin formation with accumulation of free heme (Sullivan *et al*. 1996). The detected effects on the sensor ratios might therefore be heme-mediated. To investigate further the specificity of these effects, we analyzed the interaction of CQ, as a representative of the 4-aminoquinoline group, with MQ or AQ. MQ is reported to inhibit CQ-mediated heme accumulation (Famin and Ginsburg 2002). Therefore, we predicted a preventative effect of MQ on the CQ-mediated ATeam ratio and pH drop. AQ, which belongs to the 4-aminoquinoline group, as does CQ, was used as a control in this setting. We found that, indeed, MQ but not AQ prevented CQ-mediated ATeam1.03YEMK (Fig. 17A) and sfpHluorin (Fig. 17B) ratio depletion at 100x and 10x of their EC_50_ concentrations. In contrast, AQ did not show such a preventative effect.

**Fig. 17:**
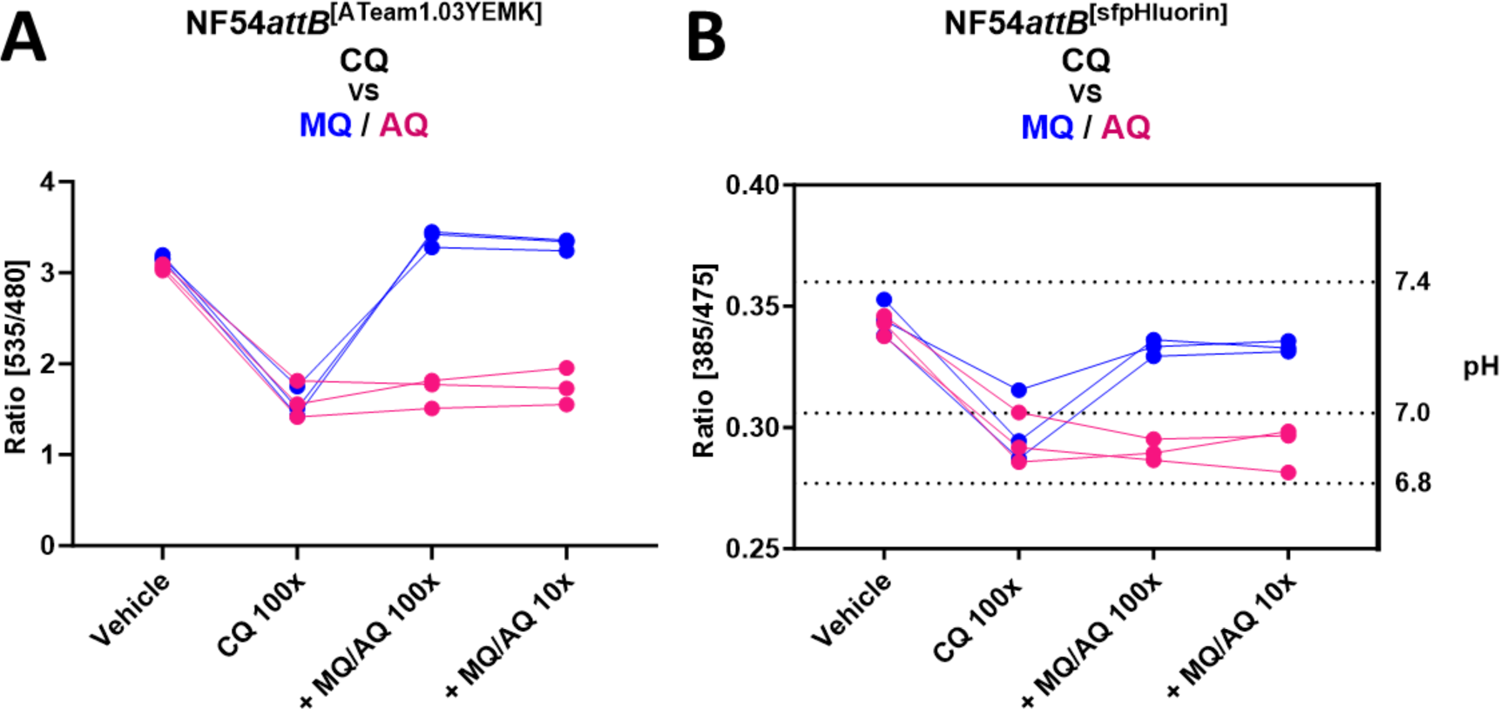
MQ but not AQ prevents CQ-mediated ATeam and sfpHluorin ratio depletion. Fluorescence microscopic measurement of MACS-enriched NF54*attB*^[ATeam1.03YEMK]^ (A) and NF54*attB*^[sfpHluorin]^ (B) trophozite-iRBCs. Each measurement corresponds to the 535 nm to 480 nm emission ratio after excitation at 430 nm, or 385 nm to 475 nm excitation ratio with emission sensing at 525, as indicated. Mean of 100 single-cell analyses after incubation with multiple of EC_50_ with n = 3 independent experiments are shown. NF54*attB*^[ATeam1.03YEMK]^ and NF54*attB*^[sfpHluorin]^ were treated either with vehicle control, 100x CQ, 100x CQ and 100x MQ, 100x CQ and MQ 10x (blue) or with vehicle, 100x CQ, 100x CQ and 100x AG, and 100x CQ and 10x AQ (magenta) of their EC_50_ concentration for 6 h.

### Mefloquine but not amodiaquine prevents chloroquine mediated ATeam sensor degradation

As shown on Fig. 13, we see a distinct emission ratio response of NF54*attB*^[ATeam1.03YEMK]^ towards 4-aminoquinoline compounds. The single-cell ratio distributions resemble a bimodal distribution with ratio allocations way below the ratio allocations of parasites starved from GLC. As the ATeam sensors are based on intramolecular FRET, such low ratios are only expected if the sensors are degraded, or, if the cytosolic pH is heavily affected. We found only moderate effects on cytosolic pH (Fig. 14). Therefore, we were interested if CQ treatment led to ATeam sensor degradation and if MQ could prevent such degradation. Thus, we carried out western blot analysis of cells treated with CQ, CQ and MQ, CQ and AQ, or vehicle control (Fig. 18).

**Fig. 18:**
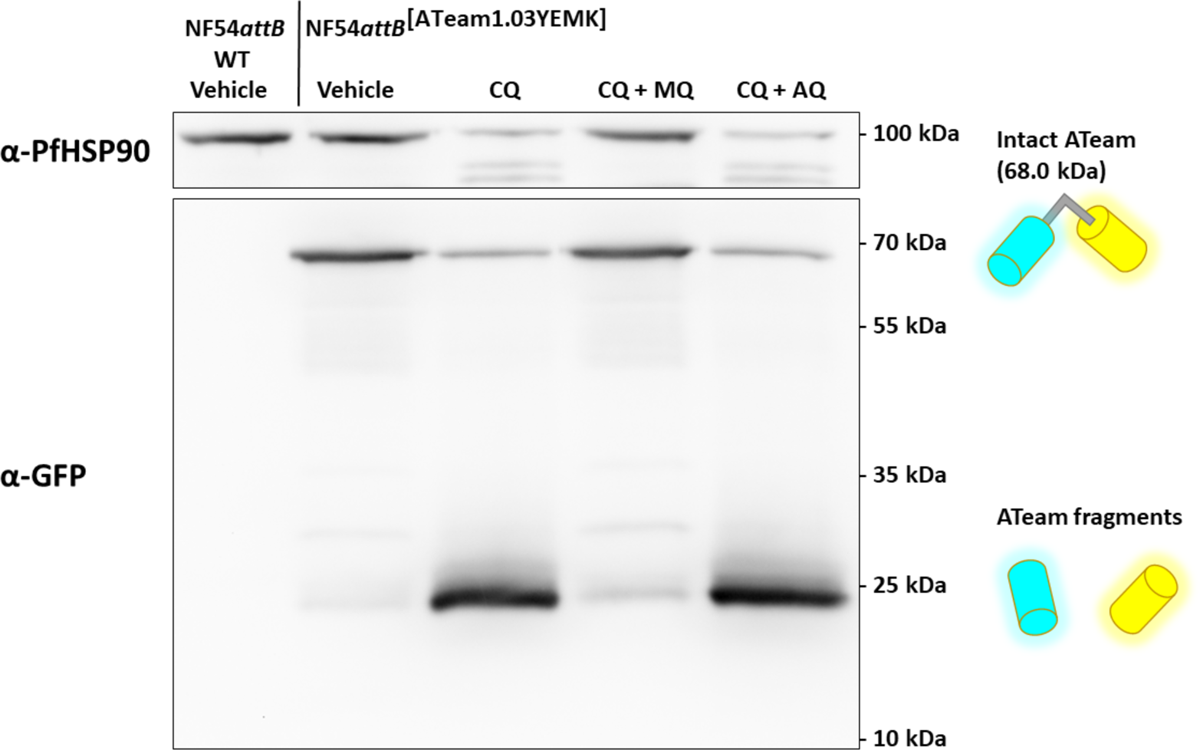
α-GFP western blot of NF54*attB*^[ATeam1.03YEMK]^ demonstrates CQ-induced sensor degradation and protective effect of MQ. α-GFP western blot of 5 million NF54*attB* (WT) or NF54*attB*^[ATeam1.03YEMK]^ trophozoite-iRBCs treated with CQ, CQ and MQ, CQ and AQ, or vehicle control at 100x EC_50_ for 6 h each.

In mock-treated NF54*attB*^[ATeam1.03YEMK]^ parasites, we could detect a clear band around 70 kDa, at the predicted size of ATeam1.03YEMK protein (68.0 kDa). Treatment with 100x aaEC_50_ CQ for 6 h caused a considerably lower band intensity at this size, but a concomitant detection of a band at around 25 kDa. This band is at the molecular weight of the protease-resistant core of GFP variants, and thus, likely indicating proteolytic cleavage of the ATeam sensor. Parasites treated with both CQ and MQ did not, or only to a marginal extent, show such a band, however, simulataneous treatment with both CQ and the 4-aminoquinoline AQ still resulted in the degradation product.

We detected PfHSP90 as a loading control. We could detect a similar PfHSP90 signal around the 100 kDa marker band in the CQ + MQ group and in the untreated control group. In the CQ and CQ + AQ treated group, we could only detect a faint signal at this molecular mass and two additional bands between 70 kDa and 100 kDa, indicating HSP90 degradation products. As housekeeping genes such as HSP90 can be up or down regulated as a stress response, or degraded under proteolytic stress, we demonstrate via Ponceau S total protein staining that the western blot signal distribution is unlikely to be explained only by unequal sample loading (Fig. S11).

## Discussion

In this report, we largely verified previous *in vitro* data for ATeam proteins (Imamura *et al*. 2009; Kotera *et al*. 2010; De Col *et al*. 2017), with small variations likely being due to slightly different experimental condition such as temperature or buffer composition. Overall, all sensors exhibited spectral properties as expected and were assessed by us as functional. We further excluded direct compound-sensor interactions as a cause of ratio changes, under the conditions studied. Our analyses did however identify a pH-dependence for ATeam1.03-nD/nA under specific conditions which must be considered when applying such sensors, especially under conditions, such as drug treatments, which may cause pH shifts. Following this, we established ATeam1.03YEMK and ATeam1.03-nD/nA measurements in *P. falciparum*. We could successfully demonstrate dynamic changes of ATP levels, both in a plate reader, as well as in live single-cell fluorescence microscopy. Both sensor cell lines exhibit spectral properties as expected and changed their emission ratio in response to GLC starvation, recovery, and subsequent 2-DG treatment in accordance with the expected corresponding ATP level. We therefore assess our measurement systems as validated. NF54*attB*^[ATeam1.03-nD/nA]^ showed a marked lower fluorescence intensity in the parasites and an unstable pH response of the recombinant protein, and was thus excluded from further analyses.

For our ATP-drug-response analyses, we are interested in qualitative changes of ATP level not absolute quantitative measurements of [ATP]. We refrained from deduction of molar [ATP] *in cellulo* from ratios using calibration of recombinant protein or *in cellulo* calibration approaches (Lerchundi *et al*. 2020; Yaginuma *et al*. 2014), as distribution of ATP species in a solution is complex (Storer and Cornish-Bowden 1976), and influence of His-tag, experimental setup, or cellular disruption cannot formally be excluded. Instead, we present the ATeam system as a reliable semi-quantitative tool for measuring ATP responses in *P. falciparum*. Based on our *in vitro* and *in cellulo* characterisation, we are certain about NF54*attB*^[ATeam1.03YEMK]^’s capabilities to measure exact cytosolic MgATP^2-^ level, even though it is difficult or impossible to extrapolate absolute molar [ATP] from our data.

As we observed a pH-dependence of the sensor under certains conditions, we established the genetically-encoded pH sensor sfpHlourin to monitor pH changes under the experimental conditions we applied. Our determinations for the resting pH ± SD of 7.34 ± 0.07 and 7.32 ± 0.12 in our plate reader and live single-cell measurement setup, respectively, are in large agreement with previously reported methods (Saliba and Kirk 1999; Hayashi *et al*. 2000; Wissing *et al*. 2002) with larger discrepancies likely due to differences in experimental conditions (Yayon *et al*. 1984; Mohring *et al*. 2017; Kuhn *et al*. 2007; Wissing *et al*. 2002). We therefore assess our sfpHluorin measurement system and our interpolated pH determinations as validated.

Having established our experimental system, we then analysed parasite responses to compounds from varying chemical classes. For our experimental setup, we used 100x and 10x EC_50_ compound concentrations to induce rapid effects in a relatively short time frame, which have been determined in previous studies to largely reflect at least the peak concentration of most of the drugs *in vivo* (Adjepon-Yamoah *et al*. 1986; Anyorigiya *et al*. 2021; Chu and Dorlo 2023; Ezzet *et al*. 1998; Gutman *et al*. 2009; Kayumba *et al*. 2008; Rijken *et al*. 2011; Walter-Sack *et al*. 2009; Newton *et al*. 2005). For setting up our drug-response assays, we used starvation from GLC as a control for total ATP depletion. Thus, ratios observed below this level would be indicative of other non-specific effects such as sensor degradation. We observed the highest ratios when treating parasites with CHX. CHX is an inhibitor of protein synthesis and as protein synthesis is the major ATP-consuming process in rapidly proliferating microorganisms (Russell and Cook 1995), its inhibition has been observed to lead to increases in cytosolic ATP concentration in other systems (Sánchez-Alcázar *et al*. 1997). Our data suggest that, following CHX treatment, ATP concentrations approach sensor saturation, indicated by the compression of ratio distribution values.

Taken individually, each parameter we measured does not allow strong conclusions to be drawn about differing modes of action. However, taken as a whole and encompassing measurement of ATP, pH and cell size, our results reveal distinct response patterns to 4-aminoquinolines, arylamino alcohols, artemisinins, a selection of redox-cyclers, as well as of DOXY, ATQ, and CHX.

Like the 4-aminoquinolines presented in this study, the arylamino alcohols QN, MQ, and LUM all inhibit formation of hemozoin *in cellulo* and increase free heme to a limited extent (Combrinck *et al*. 2013). However they also show distinct differences in their cellular effects and resistance mechanisms, as reviewed before (Fitch 2004). As resistances against CQ and MQ are inversely correlated, it is now consensus that the mode of action of arylamino alcohols, such as MQ, are distinct to that of 4-aminoquinolines and that their target most likely lies in the parasite cytosol rather than in the food vacuole (Ward *et al*. 2022). In line with this, MQ counteracts many effects of CQ such as Hb-accumulation (Famin and Ginsburg 2002), and QN inhibits CQ-induced heme-accumulation (Fitch 1998). Aside from this, and even after decades of research, the exact mechanism of action of MQ and QN is still uncertain and hotly debated (Wong *et al*. 2017; Sheridan *et al*. 2018; Dziekan *et al*. 2019), and for LUM the situation is even less clear. QN, MQ, and LUM all seem to share a downstream inhibitory effect at 10xEC_50_ on protein synthesis (Sheridan *et al*. 2018) although the exact point at which they exert their influence is unknown. Our study found increasing ATeam ratios after MQ and LUM treatment with 100x and 10x EC_50_. We found a similar trend for QN regarding its single-cell ratio distribution. With these finding we confirm Khan *et al*. (2012), who detected increased ATP level in response to 5x EC_50_ MQ. Additionally, for all three drugs we measured a reduced parasite size that could be indicative of translation inhibition. As already described for CHX, inhibition of protein synthesis seems to elevate the ATP level in *P. falciparum*. Thus, the increased ATP level following MQ or LUM treatment might be the consequence of such translational arrest.

For ART-based drugs, their modes of action are thought to be even more pleiotropic in nature than for the other compound classes. Ward *et al*. (2022) reviewed artemisinin’s mode of action, recently. Upon activation via heme iron, ART is activated to a free radical leading to alkylation of heme, proteins, lipids, DNA, and other biomolecules, and thereby, among others, interfering with Hb metabolism and inducing proteotoxic stress. In our study, we found that 10x EC_50_ of the artemisinin derivative DHA leads to an increased ATP level, while 100x EC_50_ led to decreased ATP level. Considering the large number of potential targets and biochemical pathways influenced by DHA treatment, we do not believe that it is possible to draw strong conclusions as to the exact mechanism leading to these ATP responses.

In our selection of compounds, we included two redox cyclers, PD and MB. PD is a 1,4-naphthoquinone derivative, active against all asexual stages, and a promising lead compound (Ehrhardt *et al*. 2016). In contrast, MB blue is the first synthetic chemotherapeutic that was developed. The history, redox-cycling activity, and other pleiotropic targets of MB was reviewed extensively by Schirmer *et al*. (2011). In our study, we found that both compounds exert quite similar effects, causing a moderate drop of ATeam ratio and only a slight drop in cytosolic pH. PD treatment of parasites under similar conditions to those we applied has been shown to lead to a significant shift in the cytosolic redox potential towards oxidation (Bielitza *et al*. 2015). Kehr *et al*. (2011) showed that the proteome of *P. falciparum* is highly redox-regulated including glycolytic enzymes such as PfGAPDH. One explanation of our results is that, faced with oxidative stress, the parasites upregulate the activity of the pentose phosphate pathway, at the cost of glycolysis.

DOXY and other tetracyclines are known to target the apicoplast’s ribosomal translation and kill the parasites with a delayed death phenotype (Dahl *et al*. 2006) resulting largely from inhibition of isoprenoid synthesis and subsequent loss in the second cycle of the apicoplast itself (Yeh and DeRisi 2011). At higher concentrations (up to 10 µM) first cycle effect were also noted, which could be negated by supplementation with either the isoprenoid precursor IPP, or iron, suggesting that this effect is apicoplast-dependent (Okada *et al*. 2020). Interestingly, we find that 5 µM DOXY causes an increase of cytosolic pH, while this did not occur upon addition of 1 µM, indicating that first cycle effects include misregulation of cytosolic pH, by a mechanism as yet unknown.

SBI-0797750 is a promising drug candidate with nanomolar activity targeting PfGluPho, the bi-functional fusion protein of glucose 6-phosphate dehydrogenase (G6PD) and 6-phosphogluconolactonase, and treatment of parasites at 1x EC_50_ led to a significant shift in the cytosolic redox potential towards oxidation after 4 h (Berneburg *et al*. 2022). As PfGluPho redirects glucose-6-phosphate from glycolysis to the oxidative pentose phosphate pathway, inhibition of this enzyme may be expected to allow a higher rate of glycolysis and thus higher rate of ATP production. However, we could not detect dramatic effects on the ATeam ratio, even though the single-cell ratio distribution does appear to be more compressed and elevated compared to control parasites. One explanation is that the more oxidative environment generated upon SBI treatment leads, in parallel, to a downregulation of glycolytic activity, as noted above. Alternatively, changes in ATP levels over the course of the experiment may be too small to be evident.

ATQ is an inhibitor of the ETC (Fry and Pudney 1992). Malaria parasites rely, during the blood stages, largely on energy production via glycolysis (MacRae *et al*. 2013; Sakata-Kato and Wirth 2016), and the only essential function of the ETC during these stages appears to be regeneration of ubiquinone for pyrimidine synthesis (Painter *et al*. 2007). In accordance with this, mitochondrial inhibitors are reported to cause only slight alterations of cellular ATP level over a short time frame (Fry *et al*. 1990). In our experimental setup, ATQ was the only compound that did not show any effects on the trophozoite stages after 4 or 6 hours at any of the concentrations tested. Our data fit to the observations of Wilson *et al*. (2013) that ATQ asserts its effects only on later stage parasites.

It is general consensus that the 4-aminoquinolines CQ, AQ, and PYRO kill the parasites through inhibition of hemozoin formation with induction of similar metabolomics effects (Birrell *et al*. 2020). However, they also show differences regarding the size of their effect on the distribution of heme species (Combrinck *et al*. 2013; Famin and Ginsburg 2002; Roberts *et al*. 2008). In our study, we found that all of the 4-aminoquinolines included in our study show similar distinct sensor responses. High concentrations of 100x EC_50_ led to reduced parasite size, a moderate drop of pH to at or slightly below 7, as well as ATeam ratio depletion which was indicative of a drop in [ATP]. Unusually, the single-cell ATeam ratio distribution dropped below the expected minimum ratio defined by parasites starved from GLC. One explanation for this behaviour would be sensor degradation. To investigate this phenomenon, we used CQ as a representative of the compound class, and monitored sensor integrity via western blot. Indeed, treatment of parasites with CQ led to sensor degradation. In addition to effects on ATeam sensor integrity, we also observed degradation of the PfHSP90 loading control under 100x EC_50_ CQ. Interestingly, co-treatment with MQ, which is known to block endocytosis, and thus uptake of hemoglobin (Hoppe *et al*. 2004) as well as CQ-mediated Hb accumulation (Famin and Ginsburg 2002), prevented both effects. We therefore believe the sensor degradation observed is, likely indirectly, heme-mediated, by mechanisms, which will require further investigation. Although we did not carry out western blot on parasites treated with AQ or PYRO, given the similar distinct ATeam ratio depletion, we believe the latter are highly likely to cause similar sensor degradation.

### Conclusion and future outlook

Here, we have established ATeam and sfpHluorin measurements in *P. falciparum* and demonstrated their feasibility to dynamically monitor changes of ATP and pH level in either a plate-reader or single-cell microscopy format. We show that, despite some limitations, ATeams and sfpHluorin are valuable tools for studying ATP and pH metabolism in *P. falciparum*. Using these new tools, we uncover distinct response patterns towards different compound classes, adding new momentum for mode of action studies. We believe that this system shows great potential to investigate the mode-of-action of current and future anti-malaria compounds, and thus accelerate drug development, and these tools are available on request to the malaria community. Although not herein demonstrated, we wish to note that these sensors, by virtue of their genetically encoded nature, can easily be targeted to specific subcellular compartments, and thus can be used to elucidate compartment-specific effects which may have otherwise been overlooked using conventional methods.

## Materials and Methods

### Expression and purification of recombinant ATeam and sfpHluorin protein

*E. coli* strain M15 carrying pQE-30[ATeam1.03-nD/nA] or pQE-30[ATeam1.03YEMK] were grown in lysogeny broth (LB) oriented at Bertani (1951) (1 g/L NaCl, 5 g/L yeast extract, 1 g/L tryptone) at 37 °C over night to inoculate 1 L LB to OD_600_ of 0.2 with subsequent incubation to OD_600_ of 0.6 at 37 °C. Protein expression was induced via isopropyl β-D-1-thiogalactopyranoside (IPTG) to a final concentration of 0.2 mM over night at 24 °C. Cells were collected via centrifugation at 11,448 g for 15 min at 4 °C and stored at −20 °C until purification. Cells were thawed in lysis buffer (100 mM tris-HCl, 200 mM NaCl, 10 mM imidazole, pH 8.0) enriched with 150 nM pepstatin, 4 nM cystatin, 100 μM phenylmethylsulfonylfluoride (PMSF). Cells were lysed using lysozyme and DNaseI for 30 min on ice, disrupted by sonication and collected via centrifugation at 38,000 g for 30 min. The supernatant was loaded onto a nickel nitrilotriacetic acid (Ni-NTA) column equilibrated in lysis buffer for affinity chromatography. After washing, proteins were eluted with increasing concentrations of imidazole (20 - 500 mM). Fractions containing predominantly ATeam protein were pooled and applied to a HiLoad 16/600 Superdex 200 column (GE Healthcare) equilibrated with 20 mM tris-HCl, 150 mM NaCl, pH 8.0 for size-exclusion chromatography. Fractions containing ATeam protein were pooled, and concentrated in 30 kDa cut off Vivaspin tube, supplemented with 20 % (v/v) glycerol and stored at −80 °C in single-use aliquots. sfpHluorin in pQE-30 was expressed and purified in *E. coli* M15 in the same way as ATeam protein with the following exceptions: Protein expression was induced at 37 °C; Lysis was conducted in HEPES buffer (50 mM HEPES, 300 mM NaCl, pH 7.5); Elution from Ni-NTA was conducted with 10 - 500 mM imidazole; Size-exclusion chromatography was conducted in HEPES buffer; Protein was concentrated using 10 kDa cut off Vivaspin tubes; Protein was stored at −20 °C in HEPES buffer. Concentration of proteins was either determined according to Bradford (1976) or using mVenus extinction coefficient as described by Imamura *et al*. (2009).

### Plate reader measurements of purified ATeam protein

Purified ATeam protein was mixed to a final concentration of 1 µM in ATeam buffer (100 mM HEPES, 50 mM KCl, 0.5 mM MgCl_2_, pH 7.34), except for pH response analyses, in which buffers contained 50 mM KCl, 0.5 mM MgCl_2_, and 100 mM of either MES (pH 5-6.5), HEPES (pH 7-8), or CHES (pH 8.5-9). Measurements were performed on Clariostar plate reader (BMG Labtech, Ortenberg, Germany) preheated to 37 °C. Emission ratios were calculated after background subtraction of emission at 527/10 nm and emission at 475/10 nm after excitation at 435/10 nm, respectively. Dynamic measurements over time were collected every 10 s, accordingly. Spectral scans were collected from 463 - 600 nm for each nm after excitation at 435/10 nm. Mean values of n = 3 experiments with independently produced proteins are shown.

### Plate reader measurements of purified sfpHuorin protein

Purified sfpHluorin protein was equilibrated in buffers to a final concentration of 1 µM with pH varying from 5.0-9.0 (pH 5.0 to pH 6.5: 10 mM MES-KOH, 100 mM NaCl, 5 mM EDTA; pH 7.0 to pH 8.0: 100 mM HEPES-KOH, 100 mM NaCl, 5 mM EDTA, pH 8.5 to pH 9.0: 100 mM tris-HCL, 100 mM NaCl, 5 mM EDTA) at 37 °C before use. Measurements were performed on Clariostar plate reader (BMG Labtech, Ortenberg, Germany) preheated to 37 °C. Excitation scan was recorded in the range from 340 to 490 nm with emission sensing at 530/40 nm. Mean values of n = 3 experiments with independently produced proteins are shown.

### Drug-sensor interaction studies

Purified sensor proteins were equilibrated in ATeam (100 mM HEPES, 50 mM KCl, 0.5 mM MgCl_2_, pH 7.34) or sfpHlourin (100 mM potassium phosphate, 100 mM NaCl, 0.5 mM Na_2_-EDTA, pH 7.0) buffer at 37 °C to a final concentration of 1 µM together with 10x stock solutions ranging from 100 nM to 10 µM of compound, before use. Measurements were performed on Clariostar plate reader (BMG Labtech, Ortenberg, Germany). ATeam emission ratio was calculated after background subtraction of emission at 527/10 nm and emission at 475/10 nm after excitation at 435/10 nm, respectively. sfpHluorin excitation ratio was calculated from 390/15 nm and 482/16 nm excitation and 530/20 nm emission, respectively. Mean values of n = 3 experiments with independently produced proteins are shown.

### Stable integration of sensors into NF54*attB*

We used the NF54*attB* strain lacking a selectable marker (Adjalley *et al*. 2011) for *attB-attP* recombination of *attP* plasmids, containing the sensor sequences, within the cg6 gene using the mycobacteriophage Bxb1 integrase as described elsewhere (Nkrumah *et al*. 2006). Briefly, 200 µL of 5 % iRBCs were transfected with 50 µg pDC2*attP* plasmids including blasticidin s deaminase selectable marker, conferring resistance to blasticidin S, calmodulin promotor and the respective sensor sequence, in addition to 50 µg of the integrase-encoding pINT plasmid including neomycin selectable marker, conferring resistance to geniticin (G418). For transfection, cells were mixed with plasmids and cytomix (120 mM KCl, 0.15 mM CaCl_2_, 2 mM EGTA, 5 mM MgCl_2_, 10 mM K_2_HPO_4_/KH_2_PO_4_, 25 mM HEPES, pH 7.6), transferred to a 2 mm electroporation cuvette and electroporated (310 V, 950 μF, capacitance ∞). Transfectants were selected with 2.5 µg/mL basticidin (until clonal selection) and 125 µg/mL G418 (5 days). Clonal parasite lines were generated using limiting dilution. Stable integration was verified using PCR as described by Schuh *et al*. (2018).

### P. falciparum cell culture

Parasites were cultured as described elsewhere (Trager and Jensen 1976). In brief, parasites were propagated in A^+^ red blood cells in complete media (CM) composed of RPMI 1640 medium supplemented with 0.5 % Albumax, 9 mM glucose, 0.2 mM hypoxanthine, 2.1 mM L-glutamine, 25 mM Hepes, and 22 μg/mL gentamycin at 3.3 % hematocrit and 37 °C under 3 % O_2_ and 3 % CO_2_. Cells were synchronized via 5 % sorbitol for at least 3 times before each measurement. Cells were treated with sorbitol in their developmental cycles preceding the cycle of the measurement.

### *In vitro P. falciparum* drug susceptibility assay

Half maximal effective concentration (EC_50_) of pyronaridine (PYRO) against *P. falciparum* NF54*attB* cell line were performed using SYBR Green I-based fluorescence assays according to Ekland *et al*. (2011). Serial double dilutions of the compound in CM was performed in 96-well plates (black, half area, μClear, Greiner Bio-One GmbH, Frickenhausen, Germany). Synchronized ring-stage parasites were added to each well to 0.15 % parasitemia and 1.25 % hematocrit and incubated at 37 °C for 48 h. Subsequently, SYBR Green in lysis buffer (20 mM tris-HCl, 5 mM EDTA, 0.16 % w/v saponin, and 1.6 % v/v triton X-100) was added to each well for 24 h at RT in the dark. Fluorescence was measured in a Clariostar plate reader (BMG Labtech, Ortenberg, Germany) at 494 nm excitation and 530 nm emission sensing. Curve-fitting of the growth inhibition against log compound concentration with a variable slope sigmoidal function led to the EC_50_ value.

### Trophozoite-enrichment via MACS

Briefly, sorbitol-synchronized iRBCs were transferred to MACS cell separation LD columns (Miltenyi Biotec, Bergisch Gladbach, Germany), and washed and eluted in CM. For measurements using live-cell microscopy, cells were washed and resuspended in Ringer’s solution (122.5 mM NaCl, 5.4 mM KCl, 1.2 mM CaCl_2_, 0.8 mM MgCl_2_, 11 mM D-glucose, 25 mM HEPES, 1 mM NaH_2_PO_4_, pH 7.4) to a density leading to approximately 30 parasites per field of view. For plate reader measurements, cells were counted using the improved Neubauer hemocytometer (Brand GmbH, Wertheim, Germany) and diluted to the desired parasite count.

### Western blot analysis

Western blot analysis was conducted with MACS-enriched trophozoite-iRBCs. For each sample, 5 million cells were either firstly incubated with 100x compound stock solution or vehicle control in CM, or directly washed with PBS and cOmplete Protease Inhibitor Cocktail Tablets (Roche Diagnostics GmbH, Mannheim, Germany) and lysed using M-PER^TM^ buffer (ThermoFisher Scientific, Waltham, MA, USA). Cell debris was pelleted, supernatant was incubated at 95 °C with 4x sample buffer +DTT, and separated using SDS-PAGE (sodium dodecyl sulfate - polyacrylamide gel electrophoresis) according to Laemmli (1970). Proteins were blotted on polyvinylidene difluoride (PVDF) membrane. Membrane was stained with ponceau S protein stain, washed, blocked in 5 % milk and probed with α-GFP (1:1,000 in 5 % milk; Roche Diagnostics GmbH, Mannheim, Germany), for GFP-based sensor probing, followed by secondary α-mouse IgG antibodies (1:10,000 in 5 % milk; Dianova, Hamburg, Germany). Equal loading via housekeeping genes was verified via α-aldolase or α-HSP90 western blot, after GFP-probing. All solutions for western blot staining were dissolved in tris-buffered saline with 0.05 % polysorbate 20.

### Measurements of sensor cell lines via plate reader

For plate reader measurements, 2 million NF54*attB*^[ATeam1.03YEMK]^ / NF54*attB*^[ATeam1.03-nD/nA]^ or 1 million NF54*attB*^[sfpHluorin]^ MACS-enriched trophozoite-iRBCs were washed in Ringer’s solution, mixed in 384-well small volume plates (Greiner Bio-One GmbH, Frickenhausen, Germany) and measured in Clariostar plate reader (BMG Labtech, Ortenberg, Germany) at 37 °C. For NF54*attB*^[ATeam1.03YEMK]^ and NF54*attB*^[ATeam1.03-nD/nA]^, measurement setting were chosen as described for recombinant ATeam protein earlier. For NF54*attB*^[sfpHluorin]^ spectral excitation scan was collected via emission sensing at 530/40 nm and excitation from 350 to 490 nm for every nm with 10 nm bandwidth. Ratio was calculated after background subtraction of emission at 530/20 nm and excitation at 390/15 or 482/16, respectively. Dynamic measurements over time were monitored every 10 s.

### Measurements of sensor cell lines via live-cell microscopy

For imaging, Axio Observer.Z1/7 microscope (Zeiss, Oberkochen, Germany) with plan-apochromat 63x/1.40 oil immersion DIC M27 objective and Axiocam 506 camera was used. For measurements, CM was washed off with Ringer’s solution und parasites were allowed to settle on pre-heated ibidiTreat µ-Dish 35 mm Quad microcopy dishes (ibidi GmbH, Gräfelfing, Germany) at 37 °C. Emission ratio of NF54*attB*^[ATeam1.03YEMK]^ was measured using Zeiss filter set 47 (EX BP 436/20, BS FT 455, EM BP 480/40) and 48 (EX BP 436/20, BS FT 455, EM BP 535/30) for CFP and CFP-YFP-FRET signal, respectively. Colibri 7 LED module 430 nm was used for excitation. FRET ratio equals background-corrected YFP signal at 535 nm divided by background-corrected CFP signal at 480 nm. Excitation ratio of NF54*attB*^[sfpHluorin]^ was measured using Zeiss filter set 38 HE without excitation filter (BS FT 495 (HE), EM BP 525/50 (HE)). Excitation ratio equals background-corrected GFP signal at 525 nm and excitation at 385 nm divided by background-corrected GFP signal at 525 nm and excitation at 475 nm. ROIs were defined using a semi-automatic ImageJ script. Background area was selected manually. Image analysis was conducted via ImageJ Fiji (Schindelin *et al*. 2012) 1.54f.

## Statistical analysis

Statistical analyses were performed via GraphPad Prism 10.02. Analysis of drug effects on sensor expressing cell lines was conducted via repeated measure two-way ANOVA using a general linear model for analysis of the selected randomized block design. Correction for multiple comparisons was conducted using the False Discovery Rate (FDR) with the two-staged step-up method described by Benjamini, Krieger and Yekutieli (2006). The desired FDR was set to 5 %. Calibration curves for interpolation of pH values were generated using a sigmoidal, four parameter logistic model.

## Acknowledgements

The authors gratefully thank Hiromi Imamura for providing the ATeam1.03YEMK sequence and Markus Schwarzländer for providing the ATeam1.03-nD/nA sequence. This research was funded by the German Research Foundation (grant BE1540/23-2 within the DFG Priority Program 1710 on Thiol Switches, awarded to KB) and the LOEWE Center DRUID (Project E3 awarded to KB, JMP and SR) within the Hessian Excellence Program.

## Supplemental Materials

**Fig S1:**
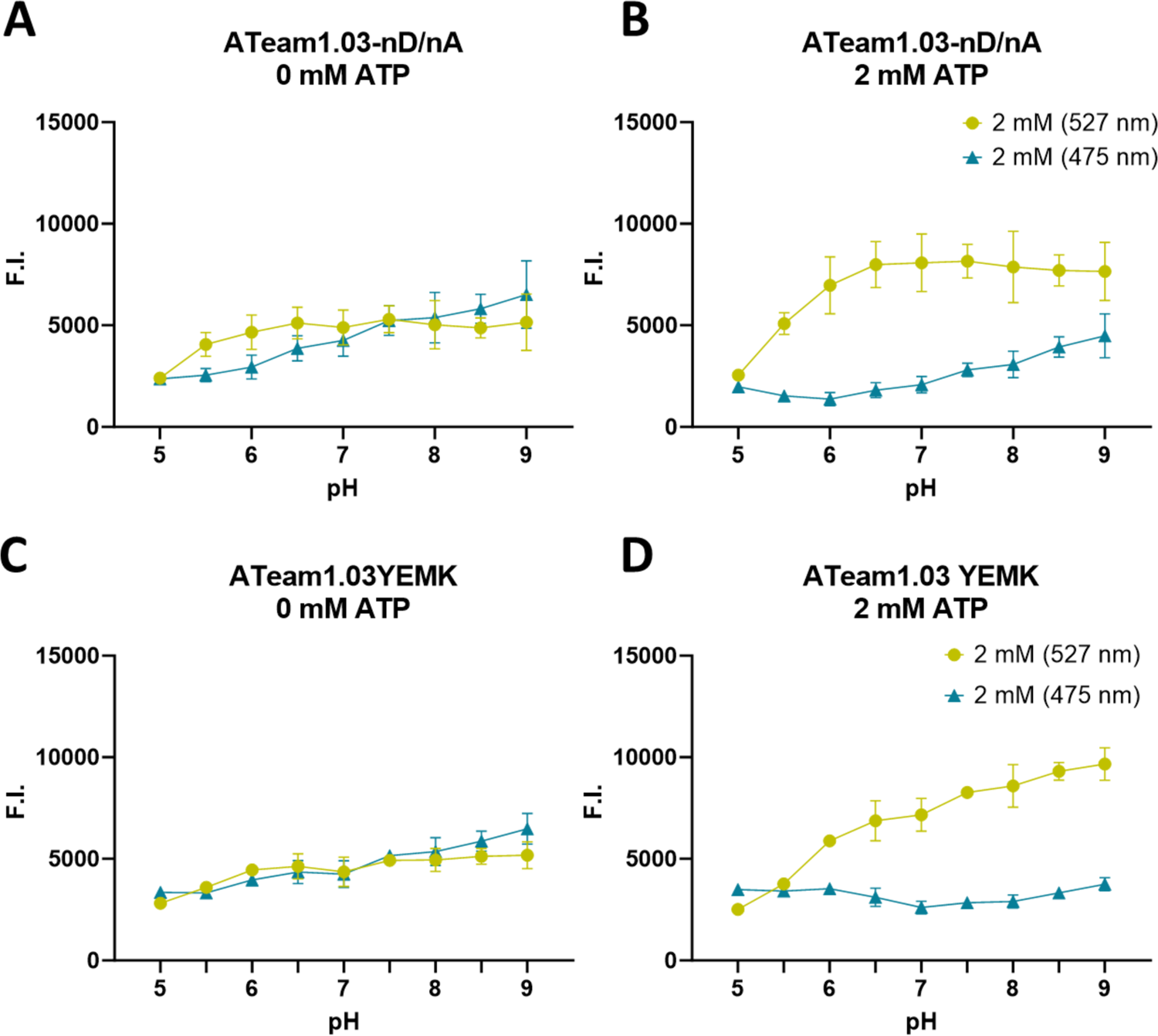
ATeam1.03-nD/nA shows less pH stability compared to ATeam1.03YEMK. A: CFP emission of ATeam1.03-nD/nA is effected by pH in the physiological range from pH 6 to 8; B: Increased dynamic range of ATeam1.03-nD/nA in response to 2 mM ATP is accompanied with decreased CFP emission; C: CFP and YFP fluorescence of ATeam1.03YEMK in the absence of ATP is stable in the physiological relevant range of pH 6 to 8; D: Ratio [527/475] decrease of ATeam1.03YEMK from pH 7 to 5 is accompanied with decreasing and increasing YFP and CFP fluorescence, respectively; Measurements were conducted via excitation at 435 nm and emission sensing at 527 nm for YFP and 475 nm for CFP. ATP was solved in equimolar solution of MgCl_2_. 1 µM recombinant protein was used. Mean values of n = 3 independent experiments are shown. Error bars indicate SD.

**Fig. S2:**
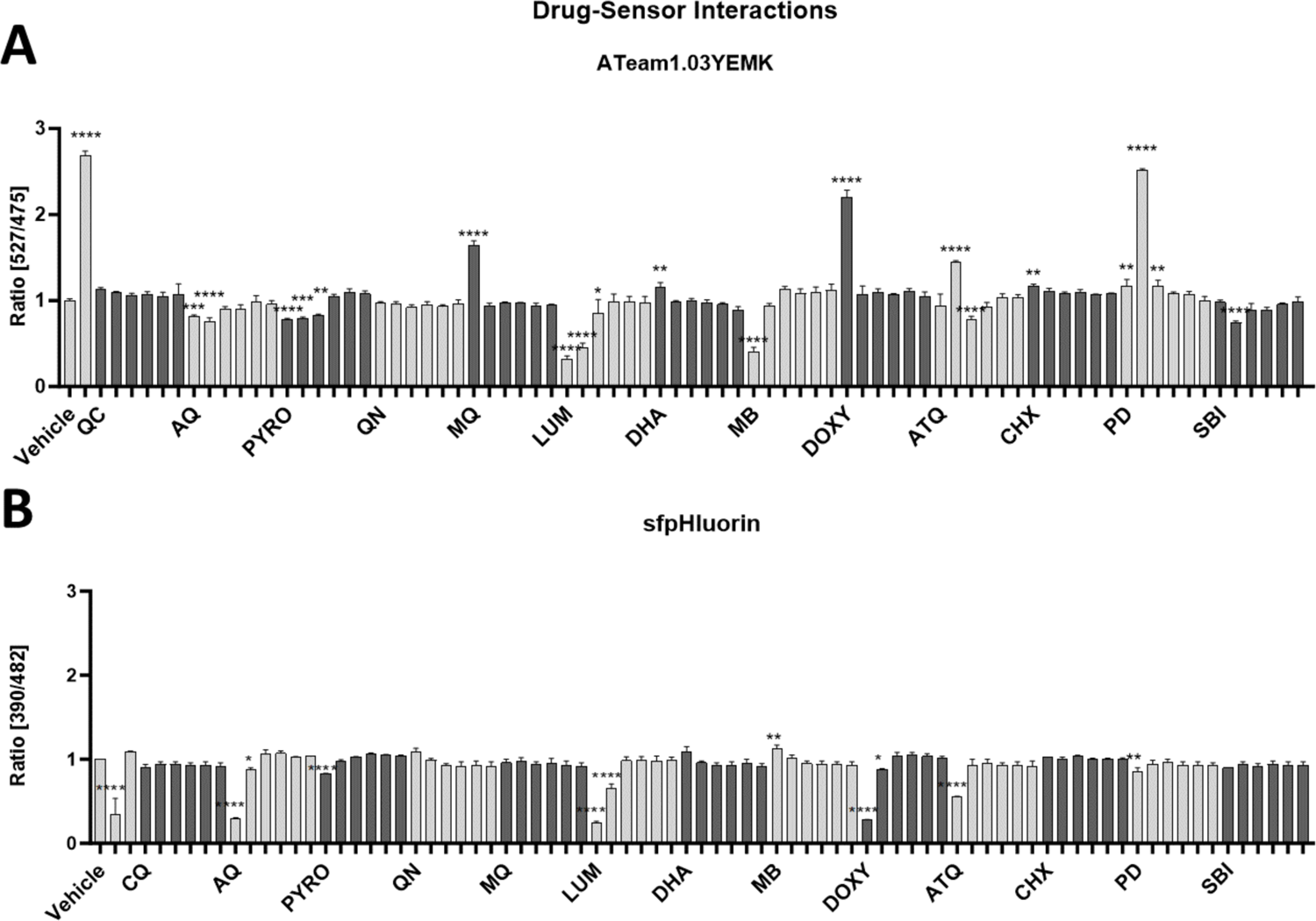
ATeam1.03YEMK and sfpHluorin drug-sensor interactions. Recombinant sensor protein was incubated with the indicated drugs (1 mM, 100 µM, 10 µM, 1 µM, 100 nM, 10 nM); A: ATeam1.03YEMK measurements included vehicle treatment with 10 mM ATP dissolved in equimolar MgCl_2_ solution; B: sfpHluorin measurements included vehicle treatment with buffers set to pH 5 and pH 9; Chloroquine (CQ), amodiaquine (AQ), pyronaridine (PYRO), quinine (QN), mefloquine (MQ), lumefantrine (LUM), dihydroartemisinin (DHA), methylene blue (MB), doxycycline (DOXY), atovaquone (ATQ), cycloheximide (CHX), plasmodione (PD), SBI 0797750 (SBI). Background-corrected emission (A) or excitation (B) ratio relative to control is shown. Sensors were incubated for 5 min. Mean ratio of n = 3 independent experiments is shown. Error bars indicate SD. Ordinary one-way ANOVA with Dunnett’s multiple comparisons testing was used. *: p < 0.05; **: p < 0.01; ***: p < 0.001; ****: p < 0.0001.

**Fig. S3:**
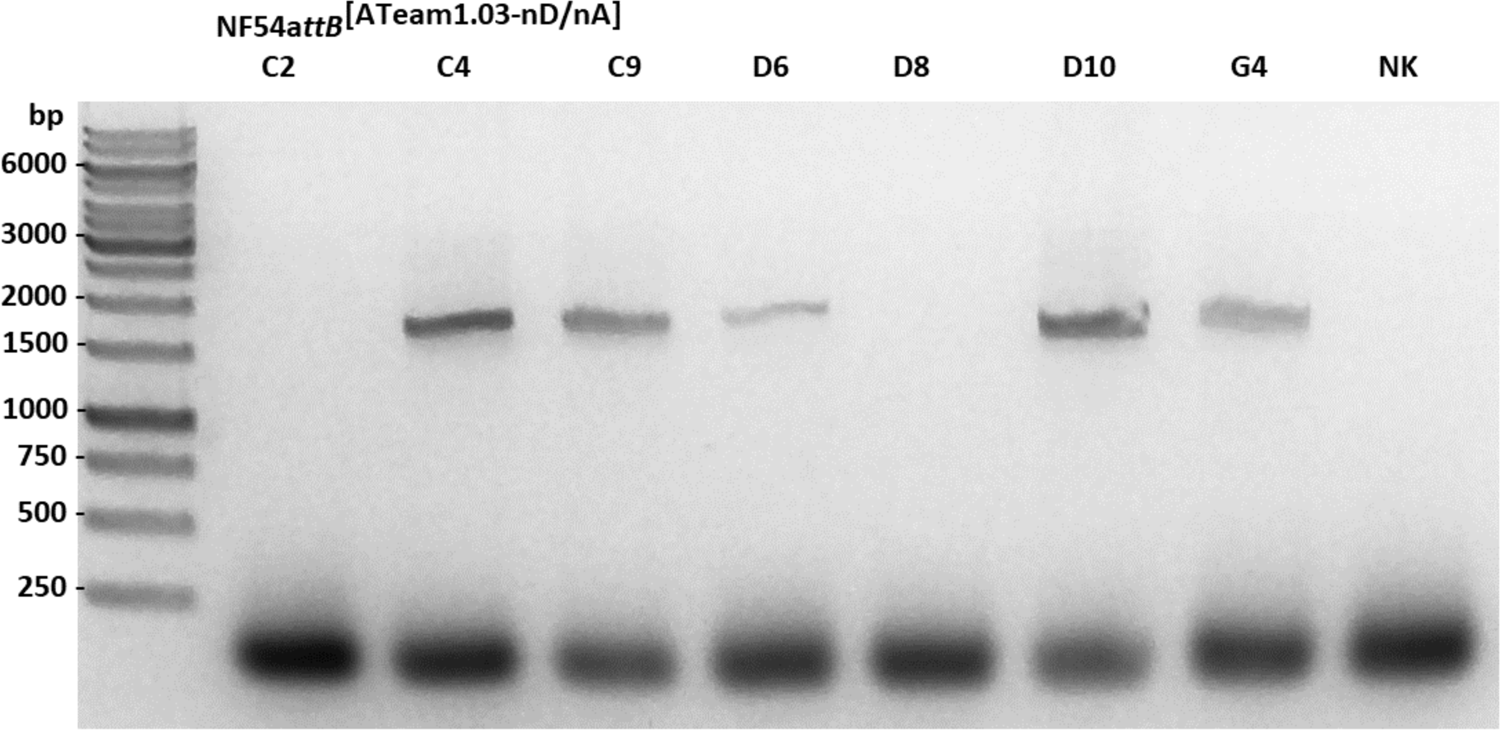
Stable integration of ATeam1.03-nD/nA in NF54*attB*. Representative clonal cell-lines show PCR product based on genomic cg6 and integrated BSD genes, and thereby, demonstrating successful sensor integration.

**Fig. S4:**
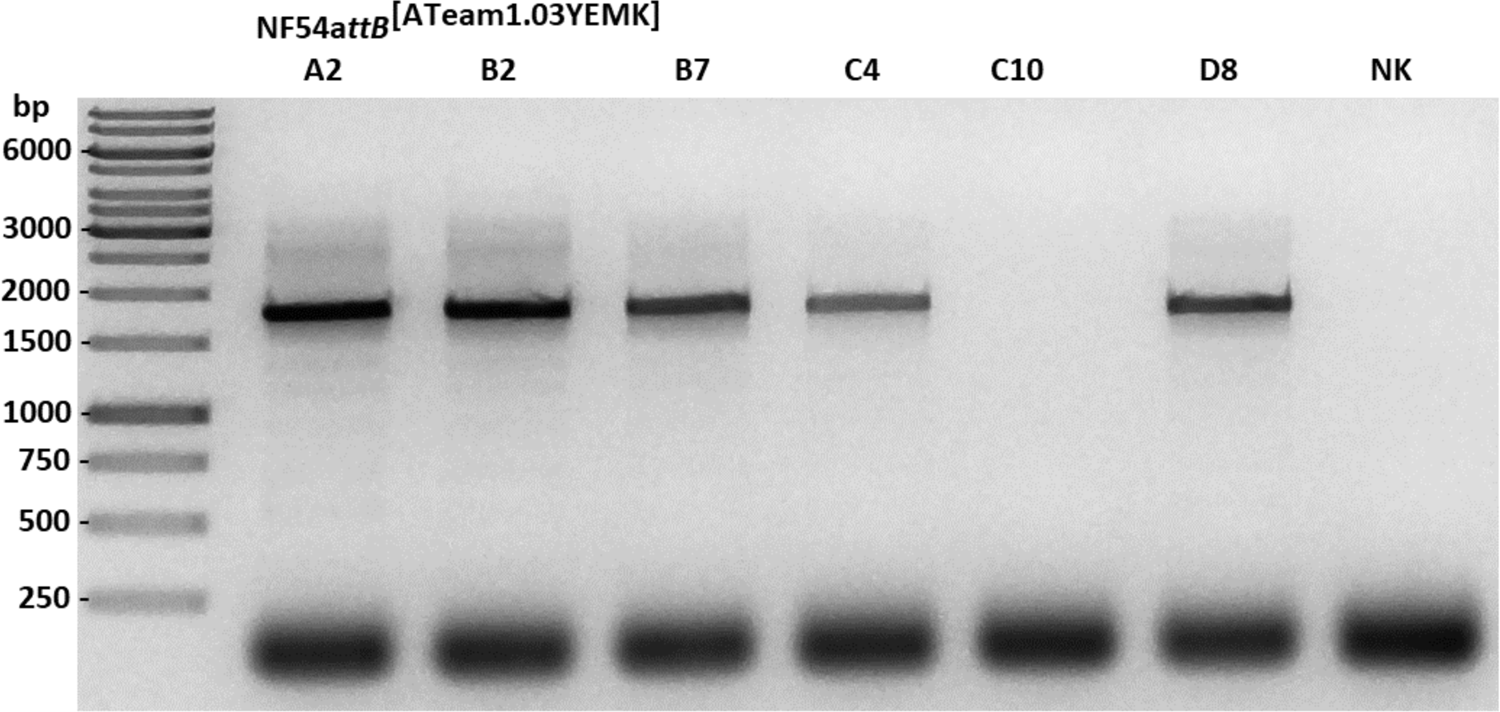
Stable integration of ATeam1.03YEMK in NF54*attB*. Representative clonal cell-lines show PCR product based on genomic cg6 and integrated BSD genes, and thereby, demonstrating successful sensor integration.

**Fig. S5:**
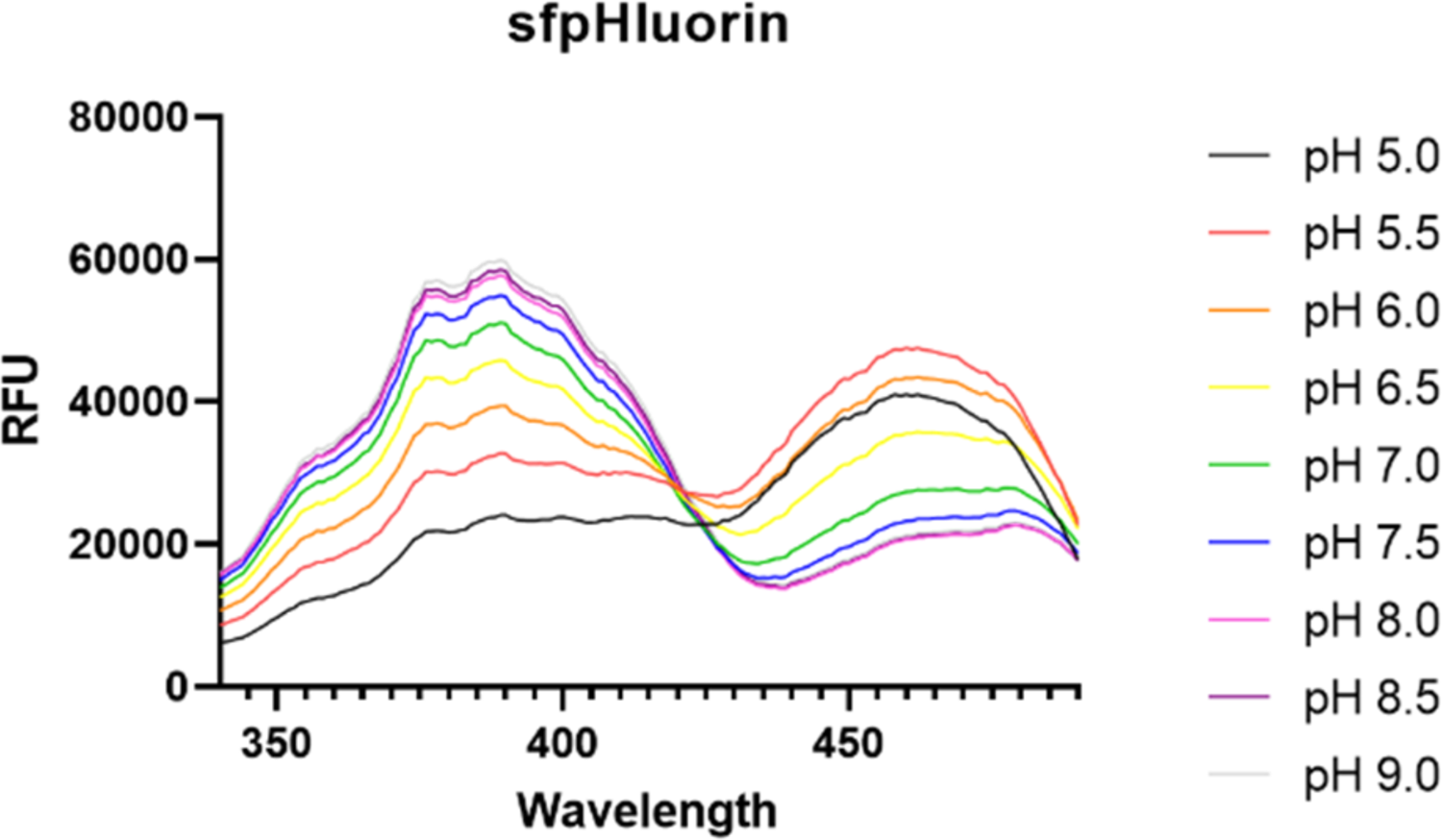
NF54*attB*^[sfpHluorin]^ *in vitro* calibration in a plate reader format. Excitation spectra from 350 to 490 nm with emission sensing at 530 nm show pH responsive peaks around 390 and 482 nm. Error bars were omitted for clarity. Means of n = 3 independent experiments are shown.

**Fig. S6:**
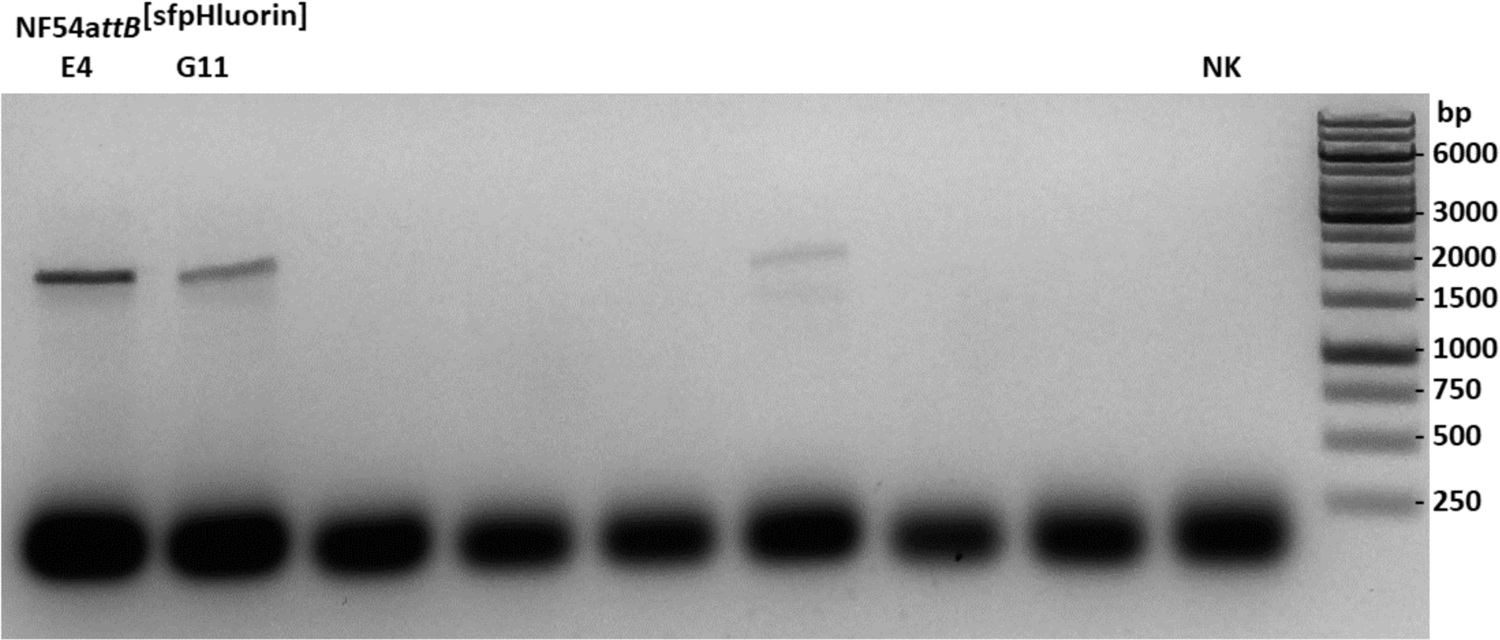
Stable integration of sfpHluorin in NF54*attB*. Representative clonal cell-lines show PCR product based on genomic cg6 and integrated BSD genes, and thereby, demonstrating successful sensor integration.

**Fig. S7:**
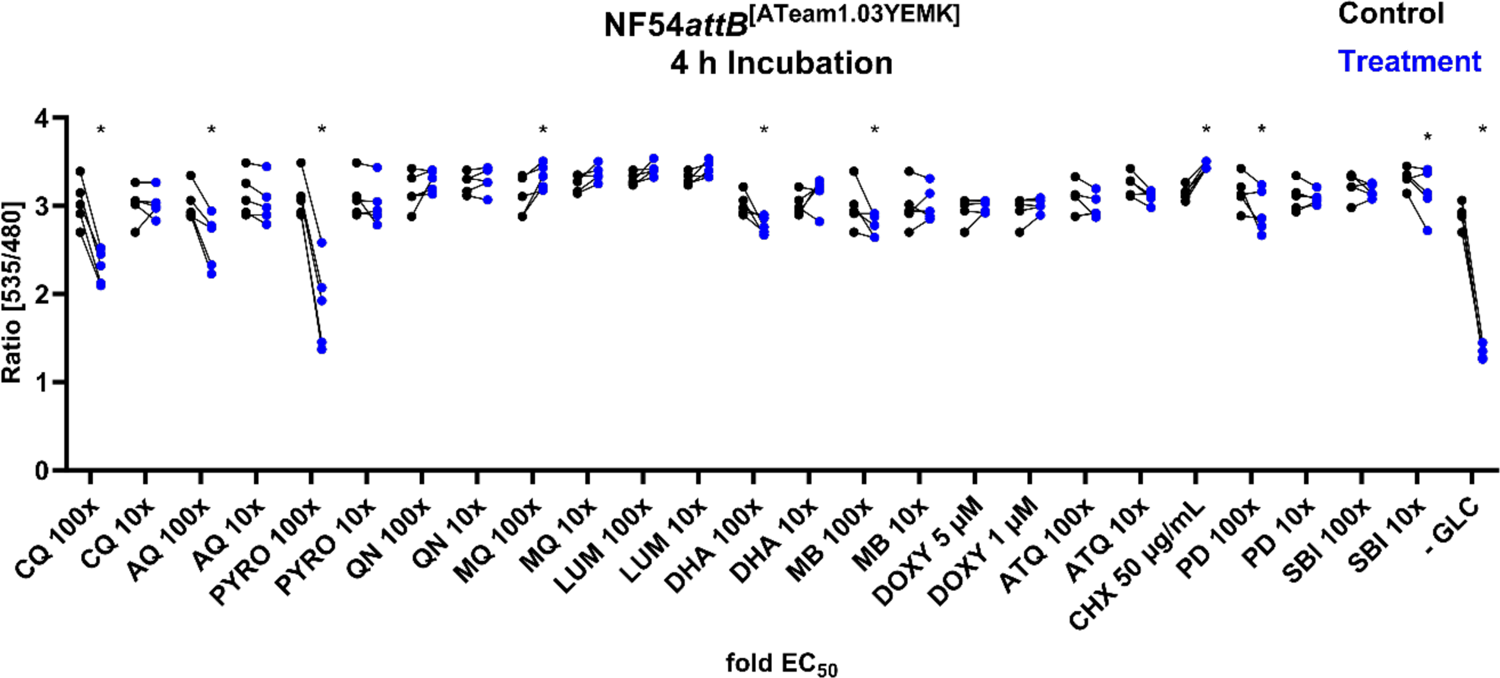
NF54*attB*^[ATeam1.03YEMK]^ shows distinct ratio changes in response towards different drug classes after 4 h incubation. Fluorescence microscopic measurement of MACS-enriched trophozite infected red blood cells. Each measurement corresponds to the 535 nm to 480 nm emission ratio after excitation at 430 nm. Mean of 100 single cell analyses of compound intervention (Treatment) and contemporaneous vehicle control (Control) with n = 5 independent experiments are shown. Treatment groups included chloroquine (CQ), amodiaquine (AQ), pyronaridine (PYRO), quinine (QN), mefloquine (MQ), lumefantrine (LUM), dihydroartemisinin (DHA), methylene blue (MB), doxycycline (DOXY), atovaquone (ATQ), cycloheximide (CHX), plasmodione (PD), SBI 0797750 (SBI), and glucose starvation (-GLC). Cells were starved from GLC through washing in PBS just before measurement. If not indicated otherwise, compounds were applied with 100x and 10x EC_50_ concentration. * indicates discovery with FDR = 5 %.

**Fig. S8:**
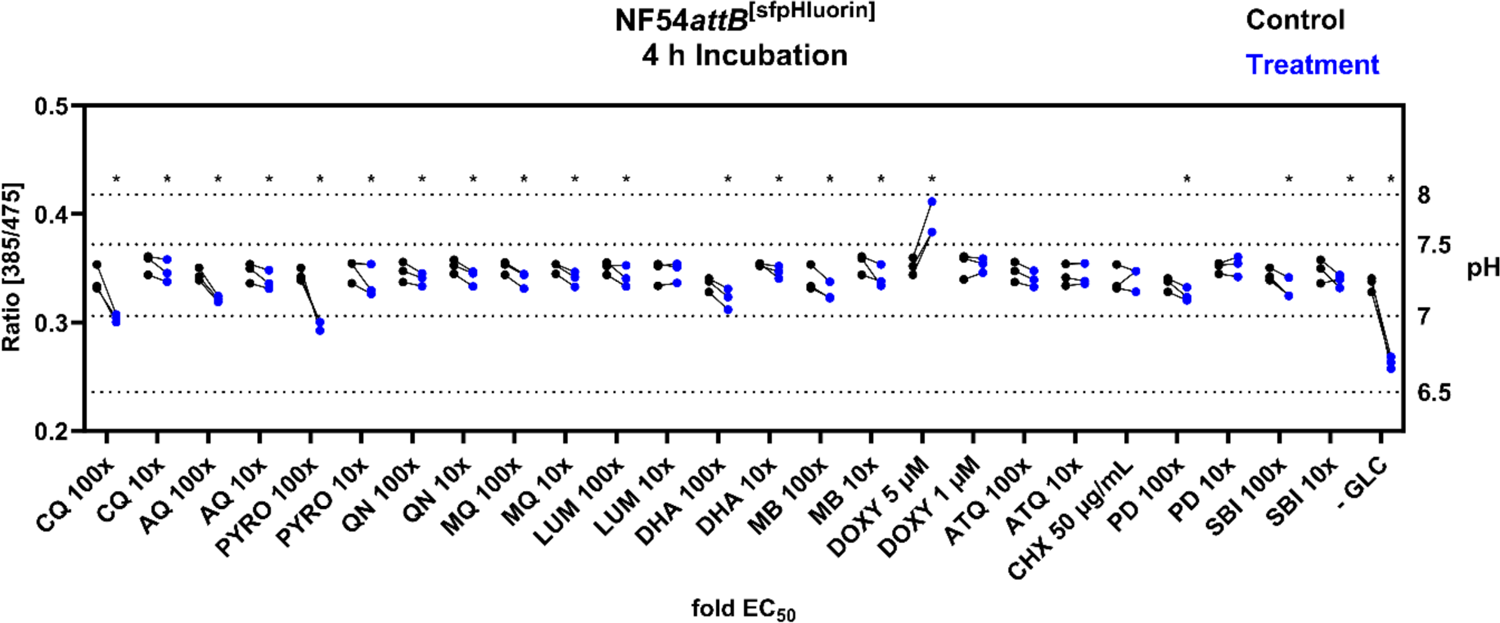
NF54*attB*^[sfpHluorin]^ shows predominantly marginal ratio changes in response towards different drug classes after 4 h incubation time. Fluorescence microscopic measurement of MACS-enriched trophozite infected red blood cells. Each measurement corresponds to the 385 to 475 nm excitation ratio at 525 nm emission. Mean of 100 single cell analyses of compound intervention (Treatment) and concurrent vehicle control (Control) with n = 3 independent experiments are shown. Treatment groups included chloroquine (CQ), amodiaquine (AQ), pyronaridine (PYRO), quinine (QN), mefloquine (MQ), lumefantrine (LUM), dihydroartemisinin (DHA), methylene blue (MB), doxycycline (DOXY), atovaquone (ATQ), cycloheximide (CHX), plasmodione (PD), SBI 0797750 (SBI), and glucose starvation (-GLC). Cells were starved from GLC through washing in PBS just before measurement. If not indicated otherwise, compounds were applied with 100x and 10x EC_50_ concentration. * indicates discovery with FDR = 5 %.

**Fig. S9:**
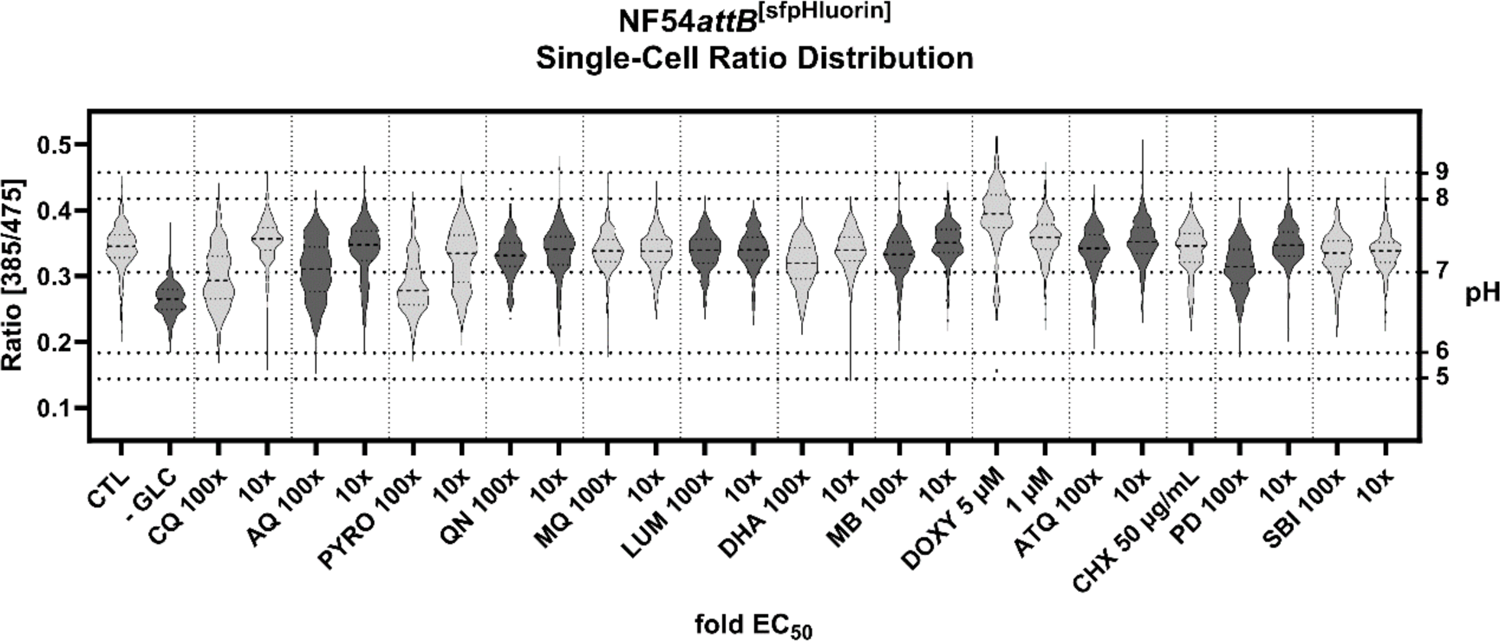
NF54*attB*^[sfpHluorin]^ shows distinct single-cell excitation ratio distribution in response towards different drug classes. Fluorescence microscopic measurement of MACS-enriched trophozite-iRBCs. Each measurement corresponds to the 385 to 475 nm excitation ratio at 525 nm emission. Pool of n = 3 independent experiments with each 100 single cell analyses is shown. Treatment groups included chloroquine (CQ), amodiaquine (AQ), pyronaridine (PYRO), quinine (QN), mefloquine (MQ), lumefantrine (LUM), dihydroartemisinin (DHA), methylene blue (MB), doxycycline (DOXY), atovaquone (ATQ), cycloheximide (CHX), plasmodione (PD), SBI 0797750 (SBI), and glucose starvation (-GLC). Cells were starved from GLC through washing in PBS just before measurement. If not indicated otherwise, compounds were applied with 100x and 10x EC_50_ concentration.

**Fig. S10.**
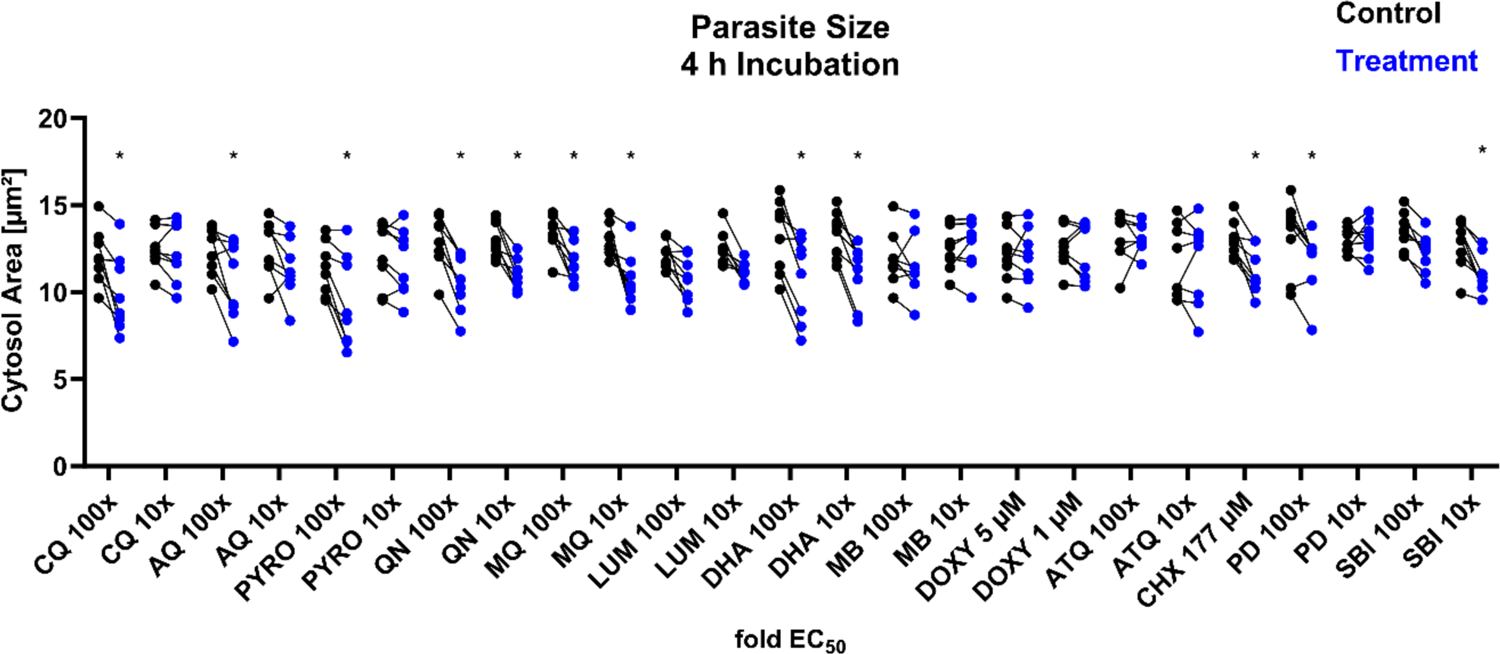
NF54*attB*^[ATeam1.03YEMK]^ and NF54*attB*^[sfpHluorin]^ show decreased parasite size in response towards different drug classes after 4 h incubation. The mean cytosol size of 100 MACS-enriched trophozite infected red blood cells derived from automated ROI definition from fluorescence microscopic measurements of NF54*attB*^[ATeam1.03YEMK]^ (n = 5) and NF54*attB*^[sfpHluorin]^ (n = 3) is combined. Analyses of compound intervention (Treatment) and concurrent vehicle control (Control) with in total n = 8 independent experiments are shown. Treatment groups included chloroquine (CQ), amodiaquine (AQ), pyronaridine (PYRO), quinine (QN), mefloquine (MQ), lumefantrine (LUM), dihydroartemisinin (DHA), methylene blue (MB), doxycycline (DOXY), atovaquone (ATQ), cycloheximide (CHX), plasmodione (PD), and SBI 0797750 (SBI). If not indicated otherwise, compounds were applied with 100x and 10x EC_50_ concentration. * indicates discovery with FDR = 5 %.

**Fig. S11:**
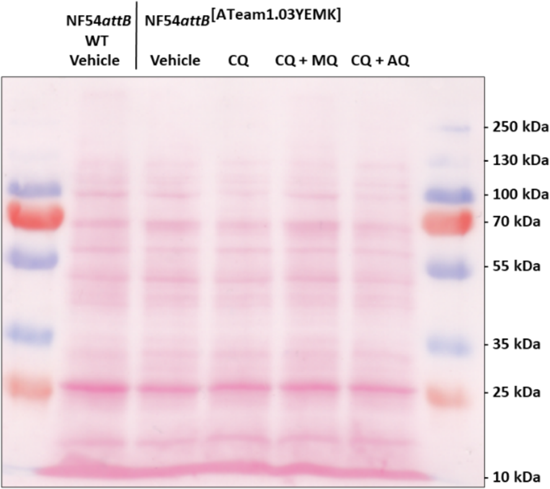
Ponceau S protein staining demonstrates sample loading.

---

**Table S1:** EC_50_ concentrations used for drug incubation of sensor cell lines. EC_50_ values were either measured (PYRO) or orientated to results in the literature.

| Compound | EC<sub>50</sub> |
| --- | --- |
| CQ | 6.9 nM (Schuh *et al.* 2018) |
| AQ | 3.4 nM (Schuh *et al.* 2018) |
| PYRO | 2.84 ± 0.77 |
| QN | 66.2 nM (Schuh *et al.* 2018) |
| MQ | 23.4 nM (Schuh *et al.* 2018) |
| LUM | 10.7 nM (Schuh *et al.* 2018) |
| DHA | 5 nM (Nardella *et al.* 2020) |
| MB | 3.3 nM (Pascual *et al.* 2011) |
| ATQ | 0.5 nM (Schuh *et al.* 2018) |
| PD | 50 nM (Cichocki *et al.* 2021) |
| SBI | 83.8 nM (Berneburg *et al.* 2022) |

**Table S2:**
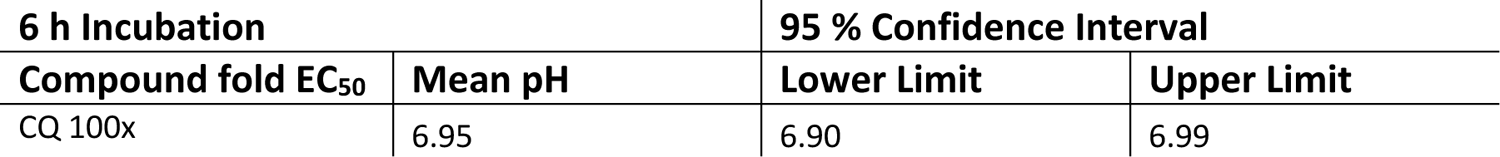
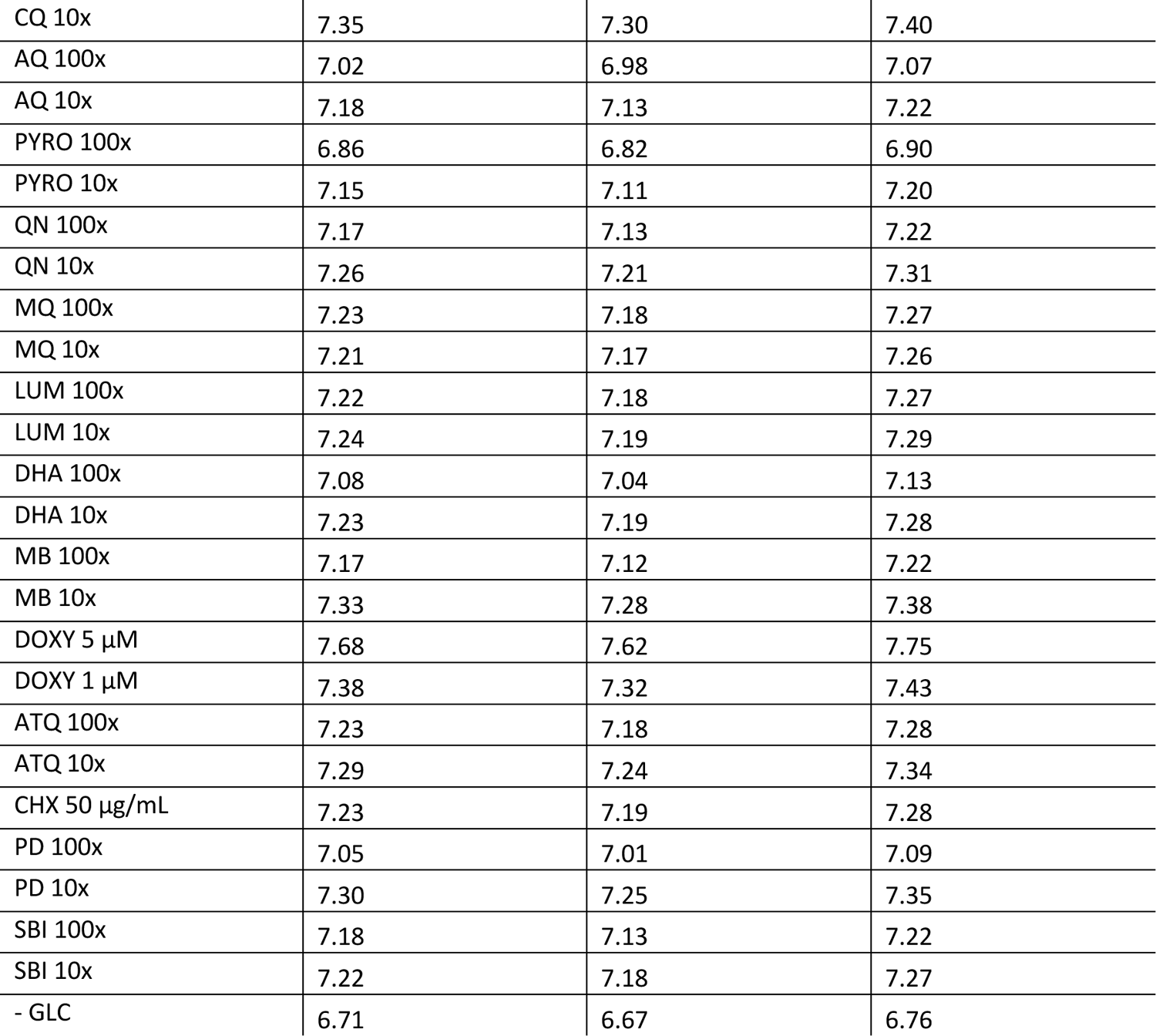
pH values of compound interventions after 6 h incubation.

**Table S3:** pH values of compound interventions after 4 h incubation.

| 4 h Incubation |  | 95 % Confidence Interval |  |
| --- | --- | --- | --- |
| Compound fold EC<sub>50</sub> | Mean pH | Lower Limit | Upper Limit |
| CQ 100x | 6.98 | 6.94 | 7.03 |
| CQ 10x | 7.29 | 7.24 | 7.35 |
| AQ 100x | 7.11 | 7.06 | 7.15 |
| AQ 10x | 7.23 | 7.18 | 7.28 |
| PYRO 100x | 6.94 | 6.90 | 6.98 |
| PYRO 10x | 7.22 | 7.17 | 7.26 |
| QN 100x | 7.24 | 7.19 | 7.29 |
| QN 10x | 7.25 | 7.21 | 7.30 |
| MQ 100x | 7.24 | 7.19 | 7.29 |
| MQ 10x | 7.24 | 7.20 | 7.29 |
| LUM 100x | 7.26 | 7.21 | 7.31 |
| LUM 10x | 7.29 | 7.25 | 7.35 |
| DHA 100x | 7.11 | 7.07 | 7.16 |
| DHA 10x | 7.29 | 7.24 | 7.34 |
| MB 100x | 7.15 | 7.11 | 7.20 |
| MB 10x | 7.25 | 7.21 | 7.30 |
| DOXY 5 µM | 7.70 | 7.63 | 7.76 |
| DOXY 1 µM | 7.34 | 7.29 | 7.39 |
| ATQ 100x | 7.24 | 7.19 | 7.29 |
| ATQ 10x | 7.26 | 7.21 | 7.31 |
| CHX 50 µg/mL | 7.25 | 7.20 | 7.30 |
| PD 100x | 7.13 | 7.09 | 7.18 |
| PD 10x | 7.33 | 7.28 | 7.39 |
| SBI 100x | 7.17 | 7.12 | 7.22 |
| SBI 10x | 7.23 | 7.18 | 7.28 |
| - GLC | 6.70 | 6.65 | 6.75 |

